# Natural variants in *C. elegans atg-5* 3’UTR uncover divergent effects of autophagy on polyglutamine aggregation in different tissues

**DOI:** 10.1101/670042

**Authors:** J Alexander-Floyd, S Haroon, M Ying, AA Entezari, C Jaeger, M Vermulst, T Gidalevitz

## Abstract

Diseases caused by protein misfolding and aggregation, in addition to cell selectivity, often exhibit variation among individuals in the age of onset, progression, and severity of disease. Genetic variation has been shown to contribute to such clinical variation. We have previously found that protein aggregation-related phenotypes in a model organism, *C. elegans*, can be modified by destabilizing polymorphisms in the genetic background and by natural genetic variation. Here, we identified a large modifier locus in a Californian wild strain of *C. elegans*, DR1350, that alters the susceptibility of the head muscle cells to polyglutamine (polyQ) aggregation, and causes an increase in overall aggregation, without changing the basal activity of the muscle proteostasis pathways known to affect polyQ aggregation. We found that the two phenotypes were genetically separable, and identified regulatory variants in a gene encoding a conserved autophagy protein ATG-5 (ATG5 in humans) as being responsible for the overall increase in aggregation. The *atg-5* gene conferred a dosage-dependent enhancement of polyQ aggregation, with DR1350-derived *atg-5* allele behaving as a hypermorph. Examination of autophagy in animals bearing the modifier locus indicated enhanced response to an autophagy-activating treatment. Because autophagy is known to be required for the clearance of polyQ aggregates, this result was surprising. Thus, we tested whether directly activating autophagy, either pharmacologically or genetically, affected the polyQ aggregation in our model. Strikingly, we found that the effect of autophagy on polyQ aggregation was tissue-dependent, such that activation of autophagy decreased polyQ aggregation in the intestine, but increased it in the muscle cells. Our data show that cryptic genetic variants in genes encoding proteostasis components, although not causing visible phenotypes under normal conditions, can have profound effects on the behavior of aggregation-prone proteins, and suggest that activation of autophagy may have divergent effects on the clearance of such proteins in different cell types.

## Introduction

Protein misfolding and aggregation underlie many human diseases, and contribute to tissue decline during aging (1, 2). In hereditary neurodegeneration, the disease-causing mutations are often directly responsible for misfolding and aggregation of the mutant protein (3, 4). For example, expansions of the CAG repeats in several different diseases lead to expanded polyglutamine (polyQ) tracts in affected proteins, which in turn result in their increased aggregation propensity (5-7). Such mutations exhibit “toxic gain-of-function” behavior, and thus a dominant, monogenic inheritance pattern. How misfolding or aggregation of these proteins cause the gain-of-function toxicity, and why they lead to disease is still incompletely understood. Two aspects of protein aggregation diseases may contribute to this difficulty. First, the behavior of mutant proteins appears to depend on the cellular environment: although they are often expressed broadly or even ubiquitously, only select subsets of cells are affected in each disease (8). The reasons for such differential susceptibility could lay in specific characteristics of the susceptible cells, such as for example the possible contribution of dopaminergic pathways in Parkinson’s disease; however, in most cases, the causes are unknown (8, 9). Second, while monogenic, these diseases show variation in the age of onset, severity, or clinical phenotypes (10). The variation is thought to result from stochastic and environmental factors, and from variants present in individual’s genetic background that act as modifiers (11-13). These genetic modifiers can affect proteins and regulatory pathways that either interact with the disease-causing mutant proteins, or are themselves impacted in disease (14). For example, modifiers of age of onset act prior to the onset of clinical manifestations and as such are expected to affect processes involved in the early pathogenic steps. Therefore, identifying natural modifier variants and their mechanisms can expand our understanding of cellular pathways involved in disease. Natural variants may also indicate pathways that differ from those found by the traditional approaches, such as association studies, mutagenesis, or RNAi screens. Importantly, because such modifiers are a part of natural genetic variation and are present in phenotypically normal individuals, they may pinpoint therapeutic routes that are less likely to cause detrimental side effects.

The most informative way to map genetic modifiers of disease is directly in human patients (13). A number of studies have shown that variants in genes other than the polyQ-expanded huntingtin (Htt) are capable of modifying the pathogenesis of Huntington’s Disease (HD) (12, 15-18). Two recent large studies of modifiers of the age of onset of HD have identified four loci on Chromosomes 3, 8 and 15 in HD subjects of European ancestry, and a locus on Chromosome 7 in a Venezuelan HD cluster (19-21). Pathway analysis implicated DNA repair pathways in European HD as having modifying effects, possibly via changing the size of the CAG repeat in somatic tissues, while the modifier locus in Venezuelan HD may act by regulating the bone morphogenetic protein (BMP) signaling. However, the causal genes and mechanisms involved are still unknown (19-21). The difficulties in using human patients in search for modifiers across aggregation diseases include the size and complexity of the human genome, the often small size of affected populations, and the possibility for the complex interactions among multiple modifiers (10, 13, 22). In addition, human studies may have limited ability to identify modifiers that are rare, or segregate in families rather than in entire affected populations. Model organisms offer a genetically tractable alternative due to the evolutionary conservation of the main cellular pathways. Expression of disease-related proteins in these organisms recapitulate many characteristics of human diseases that are related to the basic biology of protein misfolding and aggregation (23). For example, *C. elegans* and *Drosophila* models expressing polyQ-expanded Htt or ataxin-3, or isolated polyglutamine repeats, exhibit similar toxic gain-of-function behavior and the age- and polyQ-length dependent aggregation and toxicity as those seen in patients and in mammalian models (24-34). Many candidate modifying pathways identified in model organisms proved to be conserved, including insulin signaling, the heat-shock response, or regulators of proteostasis (35). Importantly, as in human disease, polyQ expansions in *C. elegans* also exhibit dependence on both the cellular environment (30, 36, 37) and the genetic background (38), despite their dominant gain-of-function behavior. We have previously shown that genetic variants coding for marginally stable proteins, although innocuous under normal conditions, can dramatically change both the aggregation and the associated toxicity of the aggregation-prone proteins, suggesting that genetic variation may directly impinge on cellular proteostasis (37, 39). Indeed, introduction of natural variation into the genetic background of polyQ-expressing animals independently modified several different aspects of polyQ behavior, including the onset and extent of aggregation, the susceptibility of different types of muscle cells to aggregation, and the resulting loss of motility and shortened lifespan (38). The polyQ aggregation in these genetically variable animals showed transgressive segregation, indicating that multiple additive or interacting alleles in parental backgrounds were acting as modifiers (38). A recent study has shown that natural variation also modulates the phenotypes and transcriptional changes caused by expression of α-synuclein transgene in the body-wall muscle cells of *C. elegans* (40). Thus, natural genetic variation within *C. elegans* wild strains can be used to investigate the mechanisms and pathways controlling the toxic effects of protein misfolding and aggregation.

Here, we dissected the genetic variation causing increased aggregation of the muscle-expressed 40-residue polyQ expansion (Q40::YFP, or Q40) in the background of a Californian wild strain of *C. elegans*, DR1350 (38). We identified a large modifier locus on Chromosome I as being causal for two phenotypes: altered susceptibility of the head muscle cells to aggregation and increased overall aggregation. These phenotypes were genetically separable, and we identified regulatory variants in a gene encoding a conserved autophagy protein ATG-5 as being responsible for the latter phenotype. The *atg-5* gene conferred a dosage-dependent enhancement of polyQ aggregation, with DR1350-derived *atg-5* allele behaving as a hypermorph. Surprisingly, animals bearing the variant *atg-5* allele showed enhanced response to an autophagy-activating drug. Because autophagy is expected to clear polyQ aggregates, we tested the effect of directly activating autophagy on the polyQ aggregation in our model, and found a striking tissue-dependence for the effect of autophagy on polyQ aggregation. Our data show that cryptic genetic variants in genes encoding proteostasis components can have profound effects on the behavior of aggregation-prone proteins, and suggest that activation of autophagy may have divergent effects on the clearance of such proteins in different cell types.

## Results

### DR1350-derived variants increase polyglutamine aggregation

We previously found that introgression of an integrated polyglutamine-encoding transgene (Q40) from the laboratory Bristol/N2 background (Q40Bristol) into the wild California isolate DR1350 resulted in strongly accelerated polyglutamine aggregation in the body-wall muscle cells, and a characteristic increase in the relative susceptibility of the normally resistant head muscle cells to polyQ aggregation (38). These two phenotypes were also present in 5 out of 21 recombinant inbred lines (RILs) derived from a cross between Q40Bristol and Q40DR1350 strains (38). To isolate the genetic variation that contributed to increased aggregation, we chose one (RIL2) that exhibited more than 2-fold increase in the number of aggregates relative to the Q40Bristol parent at the late fourth larval stage (L4) (**Fig. 1A**). We backcrossed RIL2 animals to the Q40Bristol parental strain 23 times, selecting for the F2 progeny that inherited RIL2-like phenotypes after each round of backcrossing (**Fig. 1B**). This approach ensured that the DR1350-derived RIL2 variants that contributed to the polyQ phenotypes were retained in the resulting 23x backcrossed strain, while the majority of its background was derived from the Q40Bristol parental strain. The backcrossed strain is referred to as *drxIR1*;Q40 (**Fig. 1B**). Since the increased susceptibility of the head muscles is an easy to detect qualitative phenotype that behaved in our RIL panel as a recessive trait (38), we used this phenotype during F2 progeny selection. Interestingly, the *drxIR1*;Q40 strain also retained the second polyQ phenotype - the increased overall aggregation **(Fig. 1A**), suggesting that the two phenotypes result from either linked or same natural variant(s). Age-matched *drxIR1*;Q40 animals had a higher number of polyQ40 aggregates than Q40Bristol until day 2 of adulthood, when polyQ40 aggregation reached its maximum in both strains (**Fig. 1C**). Thus, natural variants present in the wild isolate DR1350 can modify polyglutamine aggregation when introgressed into the Bristol genetic background.

**Fig 1.**
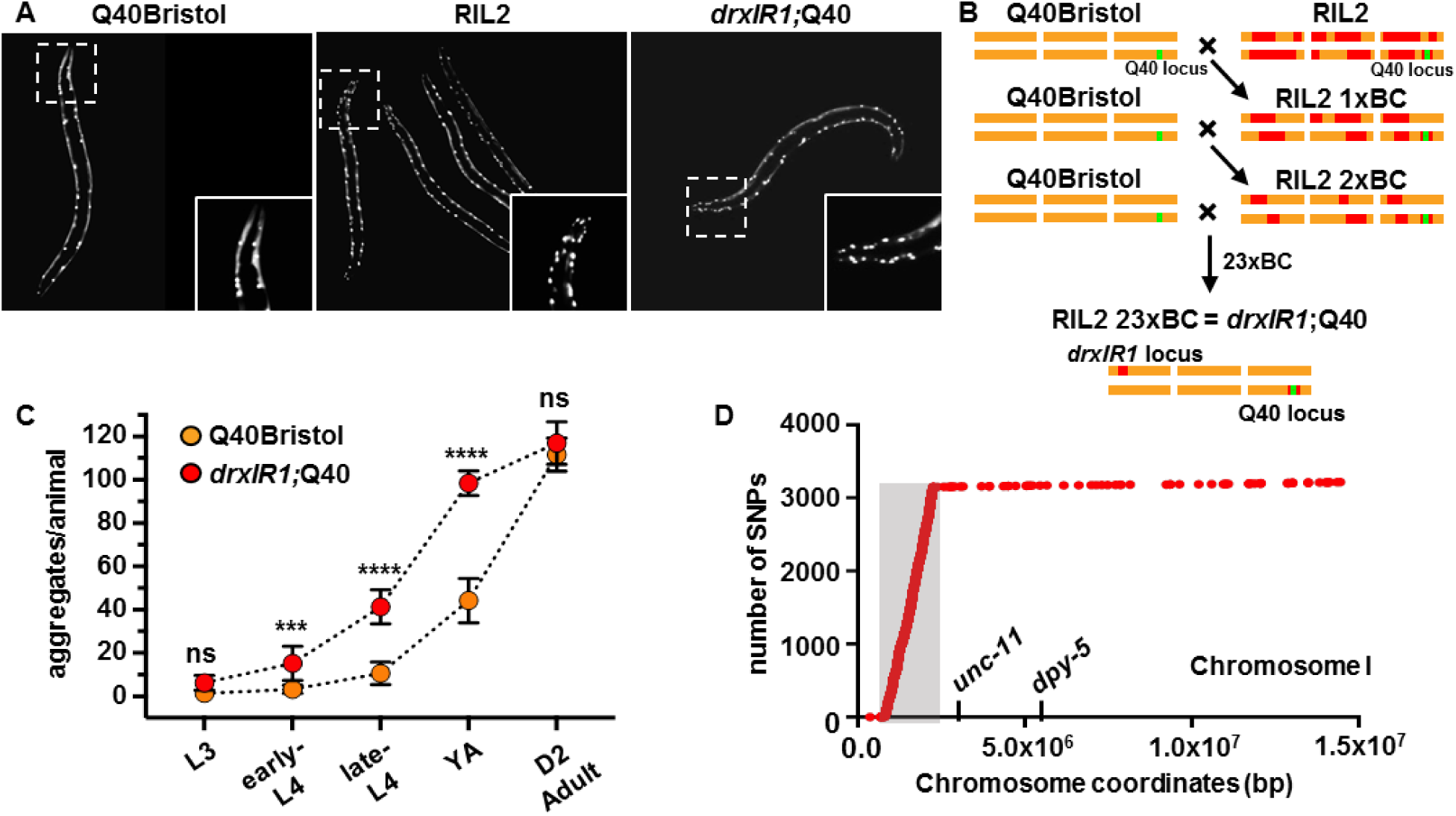
*drxIR1* locus causes increased polyQ40 aggregation. Late-L4 RIL2 and *drxlR1*;Q40 animals have increased aggregation compared to Q40Bristol animals. (**B**) The scheme for generation of the *drxIR1*;Q40 strain through rounds of crossing-selection. RIL2 strain was backcrossed (**BC**) into the Q40Bristol strain 23 times. DR1350-derived variants that are retained through the crossing-selection scheme are those contributing to the RIL2 polyQ phenotype (and their linked variants). (**C**) The *drxIR1*;Q40 animals exhibit a faster accumulation of polyQ aggregates compared to Q40Bristol at all developments stages, until both strains reach maximum at Day 2 of adulthood. L3, L4, YA and D2 adult indicate 3^rd^ and 4^th^ larval stage, young adult, and day 2 adult stage, respectively. Data are mean ± SD, 10 to 20 animals per data point. Data were analyzed by ANOVA followed by Bonferroni’s multiple comparison test, \*\*\*\**P*<0.0001,\*\*\**P*=0.0004. **Orange**: Q40Bristol background, **red**: *drxIR1*;Q40. Same color scheme is used in all figures. (**D**) DR1350-derived locus with over 3000 unique SNPs (in the grey-shaded area) to the left of *unc-11* on Chromosome I in *drxIR1*;Q40 strain.

### Polyglutamine aggregation-modifying variants reside in a large interval inherited from the DR1350 parent

In order to identify the causative variant(s) in the backcrossed *drxIR1*;Q40 strain, we first used mapping strains with visible mutations on each Chromosome, and found that increased aggregation segregated with the left arm of Chromosome I. This location was confirmed (described further below) using a free duplication sDP2 (41), which covers the left arm of Chromosome I through *dpy-5* (**Suppl. Table 1**). To precisely map the variant(s), we performed genome sequencing of both the *drxIR1*;Q40 and Q40Bristol strains and identified SNPs present only in the former, using Galaxy pipeline described in (42). We found that the left arm of Chromosome I in the backcrossed *drxIR1*;Q40 strain contained an 1.43Mb interval (ChrI:832,674-2,262,484), with over 4,000 SNPs. Because our previous data showed that introgression of the Q40 transgene into the commonly used CB4856 (Hawaiian) strain did not result in the same aggregation phenotypes as in the DR1350 background (38), we further subtracted the known Hawaiian SNPs (43) and found that the interval still contained over 3,000 SNPs (**Fig. 1D**). We tested whether this interval was also present in the remaining four high-aggregation RILs from the original study, by following several SNPs within the interval (**Suppl. Fig. 1**). We found that three of the RILs indeed inherited the entire interval, while the interval in the fourth one (RIL15) was shorter on the right side, extending through SNP 6 at ChrI:1,850,249 (WBVar00017051), but not through SNP 6b at ChrI:1,972,719 (WBVar00017376) (**Suppl. Fig. 1**). Thus, four independent RILs with high polyQ aggregation phenotypes, and the 23 times back-crossed *drxIR1*;Q40 strain derived from another RIL (RIL2), all contained the parental interval ChrI:832,674-1,972,719 from the high-aggregation DR1350;Q40 strain. To confirm, we used a mutation in *egl-30* gene located within this interval (**Suppl. Fig. 1**). We were unable to find any F2 progeny from 10 F1 heterozygotes from a cross between *drxIR1*;Q40 and *egl-30(n686)* animals that showed both the RIL2-like polyQ head aggregation phenotype and the *egl* phenotype (>1000 F2s), consistent with a close genetic linkage. Furthermore, in subsequent genetic crosses between *drxIR1*;Q40 and Q40;Bristol animals, we observed a complete correlation between F2 progeny inheriting two copies of this interval, as detected by following SNP 5 (WBVar00016276) (see Methods), and the appearance of the two polyQ phenotypes (>100 animals). Together, these data indicate that ChrI:832,674-1,972,719 interval is responsible for increased polyQ aggregation phenotypes.

The remaining part of Chromosome I contained 68 additional SNPs relative to the Q40Bristol parental strain, and all the other Chromosomes accumulated less than 200 unique SNPs each (**Suppl. Fig. 2**), consistent with previous reports (44). The large size of the modifier interval was unexpected after 23 backcrosses, suggesting that it may contain structural variants preventing recombination over this region. Alternatively, this locus could contain more than one SNP responsible for the phenotypes, perhaps distributed over the interval. Of note, the known Chromosome I *zeel-1*/*peel-1* incompatibility locus (45) was not responsible for the retention of the modifier interval through the backcrosses, as it lays outside the mapped interval (**Suppl. Fig. 1**), and does not contain DR1350-derived SNPs in the *drxIR1*;Q40 strain.

**Fig 2.**
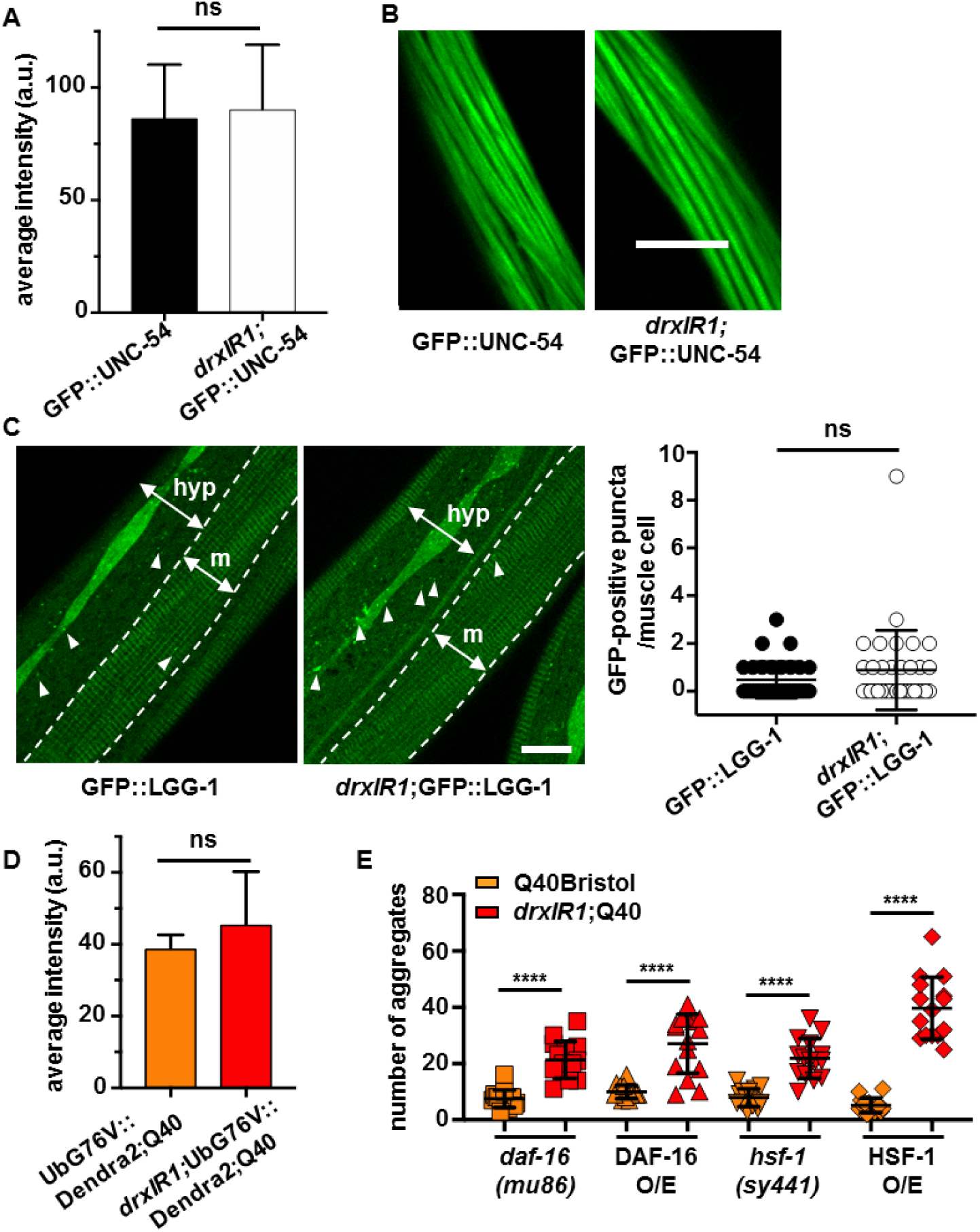
Basal protein homeostasis of muscle cells is unaffected in animals carrying the *drxIR1* interval. (**A**) Expression of GFP::UNC-54 fusion protein from *unc-54* promoter is similar between the Bristol and *drxIR1* L4 animals. Data are mean ± SD of GFP fluorescence intensity, 16-20 muscle cells per genotype, unpaired *t*-test, two-tailed. (**B**) Myofilament assembly is normal in *drxIR1* animals. Confocal images of muscle cells. Scale bar: 10 μm. (**C**) Muscle cells have very few GFP::LGG-1-positive puncta (**arrowheads**) in both Bristol and *drxIR1* L4 animals. One muscle quadrant is shown between punctate lines. **m**: muscle, **hyp**: hypodermis. An increased number of GFP::LGG-1-positive puncta is seen in the hypodermis of *drxIR1*. Scale bar is 10 μm. Right panel, quantification of GFP::LGG-1 puncta in the muscle cells. Data are mean ± SD, 30 to 40 cells (8 to 10 animals) per genotype, unpaired *t*-test, two-tailed; each symbol represents individual cells. (**D**) No difference in the average intensity of the proteasome reporter fluorescence in Q40Bristol and *drxIR1*;Q40 animals. Data are mean ± SD, 4-5 animals, unpaired *t*-test, two-tailed. (**E**) The increased aggregation phenotype in animals carrying the *drxlR1* interval does not depend on DAF-16 or HSF-1. Each symbol represents an individual animal, 15 mid-L4 animals per genotype. **O/E**: overexpression. Means ± SD are overlaid. Data were analyzed by ANOVA followed by Bonferroni’s multiple comparison test, \*\*\*\**P*<0.0001.

### Known regulators of proteostasis are not responsible for increased polyQ aggregation in *drxIR1* animals

Because the identified modifier locus contained a large number of SNPs, we thought to narrow down the candidate pathway(s) in which the modifier gene(s) acted. We first asked whether the variants in the *drxIR1* locus were increasing polyglutamine aggregation by affecting either the protein homeostasis of the muscle cells, or the Q40::YFP protein itself. We have previously tested and excluded the trivial explanation that the increased aggregation in our five RILs was due to the increased expression of the Q40::YFP protein (38). Nonetheless, we considered a possibility that *drxIR1* locus could cause increased activity of the *unc-54* promoter that was used to drive the polyglutamine transgene. To test this, we introduced an integrated *unc-54p*::GFP::UNC-54 transgene (46) into the *drxIR1* background, in the absence of polyQ, and examined its expression. We found no differences in the fluorescence levels, suggesting normal *unc-54* promoter activity (**Fig. 2A**). Since assembly of myofilaments is sensitive to both the levels of UNC-54 myosin heavy chain protein and the activity of molecular chaperones, it provides an additional measure of the GFP::UNC-54 protein levels and of the folding environment (47-49). We found normal striated pattern of GFP::UNC-54 protein in both Bristol and *drxIR1* genetic backgrounds (**Fig. 2B**).

Another reason for increased aggregation could be decreased protein turnover. To address this, we asked whether basal autophagy or proteasome activity were reduced in the muscle cells of *drxIR1* animals. Using a well-characterized autophagy reporter ubiquitously expressing GFP::LGG-1 (50), GFP::LGG-1 puncta were counted in the muscle cells of wild type and *drxIR1* animals, in the absence of the Q40::YFP protein to avoid spectral overlap. Consistent with previously published results, the number of GFP-positive puncta in the muscle cells of L4 animals with the Bristol background was low (51, 52), and we detected no difference in basal autophagy in the muscle cells of *drxIR1* animals (**Fig. 2C**), although increased number of puncta was noted in their lateral hypodermis. To test whether decreased proteasomal activity could be responsible for the increased aggregation seen in the *drxIR1*;Q40 animals, we introduced a muscle specific UbG76V::Dendra2 reporter (53) into Q40Bristol and *drxIR1;*Q40 animals, and measured its fluorescence. We detected no increase in Dendra2 fluorescence in *drxIR1* animals, indicating that there was no decrease in the proteasome activity (**Fig. 2D**). To confirm that the reporter was sensitive to decreased proteasome activity, we reduced expression of the *rpn-6.1* subunit of 19S regulatory complex of the proteasome *via* RNAi (53) and detected an increase in Dendra2 fluorescence (**Suppl. Fig. 3A**). These data indicate that increased polyglutamine aggregation in the muscle cells of *drxIR1* animals is not due to the changes in protein degradation or in polyQ protein levels.

**Fig 3.**
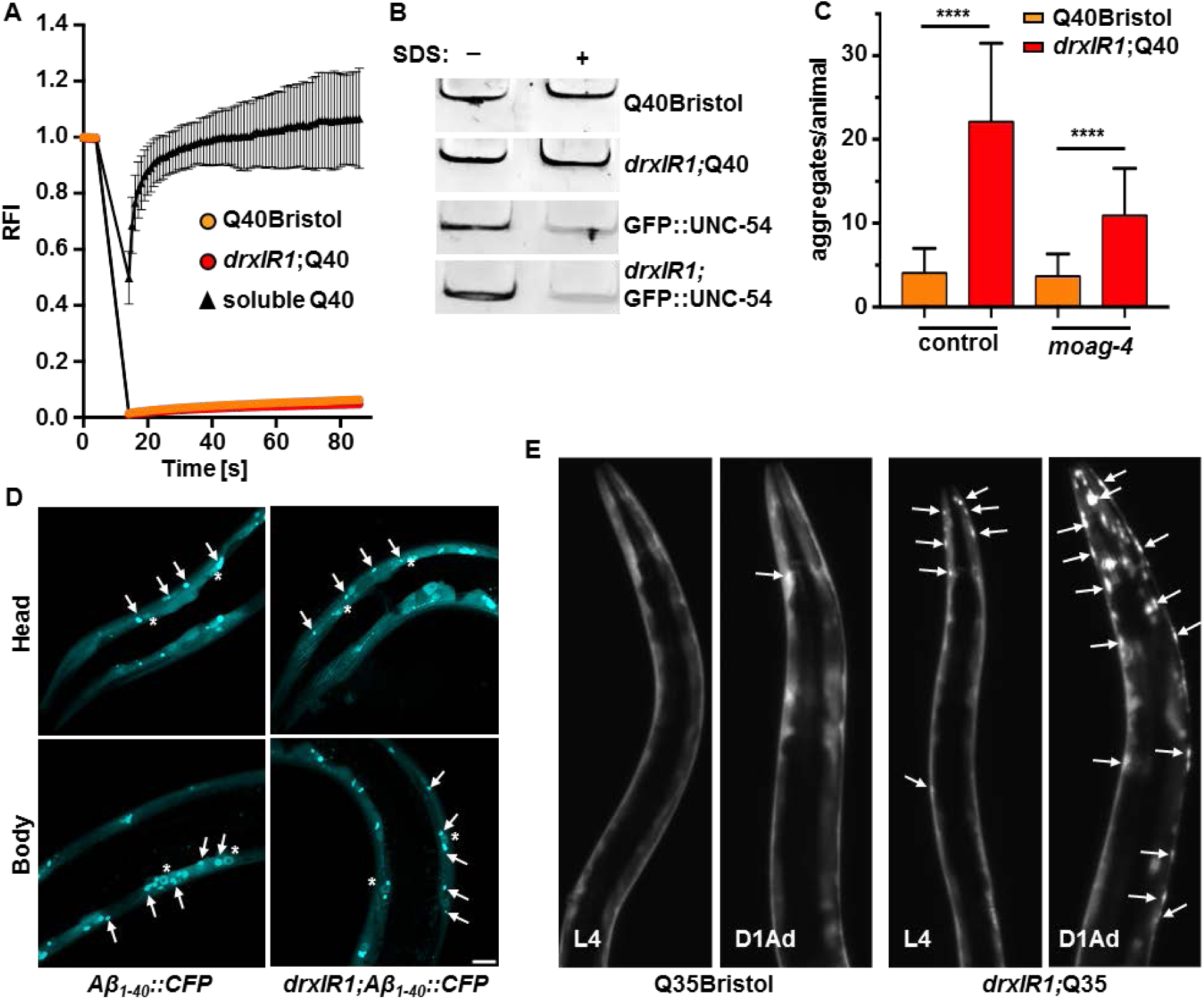
Variants in *drxIR1* interval do not alter the biophysical properties of polyQ aggregates. (**A**) FRAP analysis of fluorescent polyQ40. The soluble Q40::YFP protein recovered rapidly (**triangles**), while aggregated protein (**circles**) in both Q40Bristol and *drxIR1;*Q40 backgrounds does not recover. Data are mean ± SD. (**B**) PolyQ40 aggregates in *drxIR1*;Q40 remain SDS-resistant. Native extracts containing polyQ aggregates were treated with 5% SDS for 15 minutes and resolved by native PAGE. Aggregated proteins fail to enter the gel, remaining in the wells (shown). Native extracts containing the fibrillar GFP::UNC-54 protein were used as controls. (**C**) The increased aggregation phenotype in animals carrying the *drxlR1* interval does not depend on the amyloid-specific modifier *moag-4. moag-4* RNAi decreased the total number of aggregates in both backgrounds (YA animals are shown in Suppl. Fig. 3B), but preserved the increased aggregation in *drxIR1*;Q40 animals relative to Q40Bristol. Data are mean ± SD, three independent experiments. Data were analyzed by ANOVA followed by Bonferroni’s multiple comparison test, \*\*\*\**P*<0.0001. A total of 38 to 46 mid-L4 animals per condition. (**D**) Aggregation of a different amyloid protein, Aβ_1-40_::CFP, in unaffected by the *drxlR1* locus. Shown are confocal stacks, **arrows** point to aggregates, **asterisks** indicate Aβ_1-40_::CFP accumulating in the nuclei of the muscle cells. Scale bar: 10μm. (**E**) The shorter polyQ expansion (Q35::YFP) exhibits both the increased susceptibility of the head muscle cells and the accelerated overall aggregation in animals carrying the *drxlR1* interval. Shown are stereo micrographs, **arrows** point to some of the aggregates. **D1Ad**: day 1 adults.

**N**ext, we tested two main transcriptional pathways known to regulate cytosolic protein homeostasis - insulin signaling and the heat-shock response. Increased activity of DAF-16/FOXO, the transcription factor of the insulin signaling pathway, is associated with improved proteostasis and has been shown to affect polyglutamine aggregation (30, 36). We found that neither genetic inactivation of *daf-16*, using *daf-16(mu86)* mutation (54), nor overexpression of active DAF-16::GFP protein (55), were able to revert the increased aggregation seen in *drxIR1;*Q40 animals (**Fig. 2E**). HSF-1/HSF1 is the heat shock transcription factor that functions as a master regulator of molecular chaperones, degradation machinery, and other proteostasis components in the cytosol, and has also been shown to affect polyQ aggregation in wild type animals (36). Similarly to DAF-16, neither the hypomorphic *hsf-1(sy441)* allele, deficient in the heat-shock response (56), nor HSF-1 overexpression (57), were able to revert the increased aggregation caused by *drxIR1* background (**Fig. 2E**). Together, these data indicate that the DR1350-derived variants in *drxIR1* are not likely to act by modifying the basal proteostasis of the muscle cells of *C. elegans*.

### Variants in the introgressed interval do not alter biophysical properties of polyQ40 aggregates

Besides changes in the cellular proteostasis of the muscle cells, the increased aggregation in *drxIR1*;Q40 animals could reflect changes in the amyloid-like nature and/or biophysical properties of polyQ40 aggregates themselves. PolyQ40 is known to form immobile aggregates that do not recover after photobleaching, and are resistant to treatment with the detergent SDS (30, 58). Thus, we tested whether the presence of *drxIR1* interval altered these properties of polyQ40 aggregates. As expected, photobleaching foci within Q40Bristol resulted in essentially no recovery of fluorescence, while soluble Q40::YFP protein rapidly recovered to pre-bleach levels (**Fig. 3A**). We found no difference in recovery of Q40::YFP foci between *drxIR1*;Q40 and Q40Bristol animals (**Fig. 3A**), indicating similarly immobile aggregates. To test for SDS resistance, we extracted aggregates from Q40Bristol and *drxIR1*;Q40 animals and treated them with 5% SDS at room temperature, as described in (39). We found polyQ aggregates to be similarly SDS resistant in both genetic backgrounds (**Fig. 3B**). To confirm that our SDS treatment could dissociate non-amyloid protein assemblies, we tested GFP::UNC-54 protein that forms myofilaments (as shown in Fig. 2B). Filamentous GFP::UNC-54 protein was efficiently dissociated by SDS treatment in extracts from both Bristol and *drxIR1* backgrounds (**Fig. 3B**). recently discovered positive regulator of aggregation, MOAG-4/SERF, which specifically distinguishes amyloid and non-amyloid aggregation (59, 60), was shown to affect Q40::YFP protein in *C. elegans*. Decrease of *moag-4* expression *via* RNAi suppresses aggregation of polyglutamine, amyloid-beta (Aβ), and α-synuclein, but not of mutant SOD1 (60). To test whether the variants in the *drxIR1* background act through MOAG-4, expression of *moag-4* was knocked down by RNAi in Q40Bristol and *drxIR1*;Q40 animals. *moag-4* RNAi strongly decreased polyQ40 aggregation in both backgrounds, confirming the amyloid-like nature of aggregation in both (**Fig. 3C** (L4 animals) and **Suppl. Fig. 3B** (young adults)). However, *drxIR1*;Q40*;moag-4*(RNAi) animals retained higher aggregation relative to Q40Bristol*;moag-4*(RNAi) animals (**Fig. 3C**), as well as the increased susceptibility of the head muscles (**Suppl. Fig. 3B**), arguing against the *drxIR1* interval variants acting through MOAG-4-mediated mechanism. Together our data suggest that neither decrease in muscle proteostasis nor changes in the aggregation pathway are responsible for the increased aggregation in *drxIR1*;Q40 animals.

### The increased aggregation is specific to polyglutamine expansions

To determine whether the variants responsible for increasing polyQ40 aggregation in *drxIR1*;Q40 animals were acting generically on any amyloid aggregates, we asked if they can modify an aggregation-prone Aβ peptide. We chose the muscle specific Aβ_1-40_::CFP transgene because it exhibits both soluble and aggregated protein early in adulthood, allowing us to detect the potential increase in aggregation. We found that introduction of the *drxIR1* interval did not increase Aβ aggregation (**Fig. 3D**). In contrast, when the *drxIR1* locus was introduced into another polyglutamine model, Q35Bristol, we observed both the overall increase in polyQ35 aggregation and the increased susceptibility of the head muscles (**Fig. 3E**).

These data indicate that the DR1350-derived variants in *drxIR1* background act by a polyglutamine-specific mechanism that is likely distinct from the known aggregation-modifying mechanisms. In addition, the effect on the Q35::YFP and Q40::YFP but not on Aβ_1-40_::CFP transgenic proteins confirms that the novel mechanism acts at the protein level, rather than by modifying the transgene genomic environment, since all three transgenes were made by the same approach.

### Increased polyQ40 aggregation in the body-wall muscle cells and increased susceptibility of the head muscles to aggregation are caused by genetically separable mechanisms

Since we were unable to narrow down the candidate genes by identifying affected pathways, and our data pointed to a potentially novel pathway, we turned to an unbiased investigation of genes in the interval. As we previously reported (38), the increased susceptibility of the head muscles to aggregation (RIL2-like phenotype, measured as the ratio of head to body aggregation) behaves as a recessive trait (**Suppl. Table 1, top row**), and is fully suppressed in *drxIR1* heterozygous (*drxIR1/+*;Q40) animals. Thus we asked whether it was caused by a loss-of-function of a gene or genes in the interval, by testing whether it can be rescued in the *drxIR1* homozygotes by introducing a wild-type copy of the interval. We used a free duplication sDp2 that covers the left arm of Chromosome I, through *dpy-5* gene in the center of the Chromosome (41). Introduction of sDP2 into animals homozygous for the *drxIR1* interval and for the known loss-of-function *dpy-5(e61)* allele suppressed both the *dpy* and the RIL2-like head phenotypes to the same extent (**Suppl. Table 2, second row**), indicating that the head-muscles susceptibility phenotype in *drxIR1* animals is caused by a loss-of-function variant(s), and therefore can potentially be identified by RNAi approach in Q40Bristol animals.

In contrast, the second polyQ phenotype, the increased overall aggregation (as scored in the body-wall muscles alone, excluding the head muscles), was not suppressed in animals heterozygous for the *drxIR1* interval (**Fig. 4A**). Moreover, introduction of the sDP2 duplication, carrying the wild-type (Bristol) copy of this interval, into either Q40Bristol or *drxIR1*;Q40 animals resulted in sharply increased aggregation of polyQ40 in the body-wall muscles, relative to the corresponding strains without the duplication (**Fig. 4A**). This suggests that the phenotype of increased aggregation in the body-wall muscles depends on the dosage of a gene or genes within the boundaries of the modifier interval, and that in *drxIR1*;Q40 animals this gene carries hypermorphic variant(s), mimicking increased gene dosage. Thus, the candidate gene may be identified by RNAi approach in *drxIR1*;Q40 animals.

**Fig 4.**
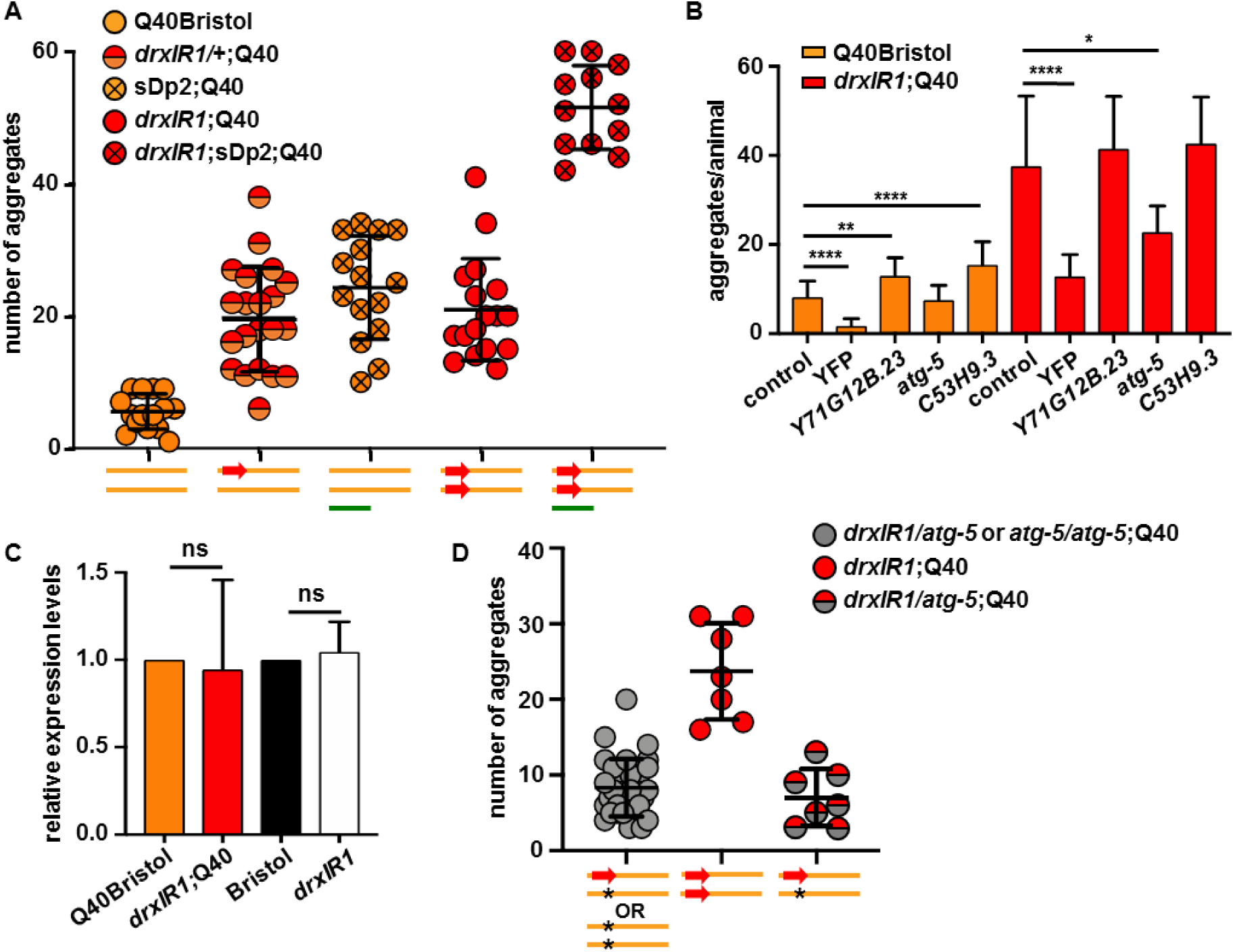
Hypermorphic variants in the autophagy gene *atg-5* are responsible for the increased polyQ aggregation in the body-wall muscles. PolyQ aggregation in the body-wall muscles is sensitive to the dosage of the *drxlR1* interval, with DR1350-derived interval acting as a hypermorph relative to the Bristol-derived interval. Each symbol represents an individual mid-L4 animal; overlaid are means ± SD. Schematic under the graph represents the genetic composition of Chromosome I: Bristol background (**orange bar**), DR1350-derived *drxlR1* interval (**red arrow**), and the free duplication sDp2 (**green bar**). (**B**) RNAi of three candidate genes affects polyQ40 aggregation. *atg-5* RNAi suppresses the increased polyQ aggregation in the muscle cells of *drxlR1* but not in Q40Bristol animals. RNAi against YFP downregulates expression of Q40::YFP protein. Data are mean ± SD, three independent experiments, 9 to 15 animals per experiment per genotype. Data were analyzed by ANOVA followed by Bonferroni’s multiple comparisons test, \*\*\*\**P*<0.0001, \*\**P*=0.0029, \**P*=0.0125. (**C**) Relative expression of *atg-5* mRNA is unaffected by the DR1350-derived *drxIR1* interval. Three independent experiments, statistics as in panel B. (**D**) *atg-5(bp484)* loss-of-function allele reverses increased aggregation caused by one copy of the DR1350-derived *drxIR1* interval. Schematic under the graph as in panel A, **star**: *atg-5* mutation. Animals were scored at mid-L4 as in panel A, compare *drxIR1/+*;Q40 animals (red/orange symbols) in panel A with *drxIR1/atg-5*;Q40 animals (red/grey symbols) in panel D. **Grey symbols** represent animals that were assumed (but not confirmed) to be heterozygous for the *drxIR1* interval, because they did not show the RIL2-like phenotype head muscle phenotype and because *atg-5/atg-5* animals exhibit strong developmental delay. Heterozygosity of *drxIR1/atg-5*;Q40 animals (**red/grey symbols**) was confirmed by singling them out and scoring segregation of the RIL2-like phenotype among their progeny. Each symbol represents an individual animals, overlaid are means ± SD.

### Autophagy-related gene 5 (ATG-5) is responsible for increased aggregation

To decrease the number of genes that were to be tested by RNAi, we were able to further narrow the large *drxIR1* interval to approximately 326 Kb (ChrI:1,647,221-1,972,719) by additionally backcrossing the *drxIR1*;Q40 animals and using the SNPs in the interval to detect recombination. The smaller 326 Kb interval contained 57 total genes including 25 candidate protein-coding genes with potentially functionally-significant SNPs (based on SnpEff annotations (62), see Methods), with 24 candidate genes remaining after exclusion of *egl-30* (**Suppl. Table 2 and Data File 1**). Each of the candidate genes was knocked down by feeding RNAi in both Q40Bristol and *drxIR1*;Q40 animals, followed by quantification of polyQ aggregation.

None of the RNAi clones affected the increased susceptibility of the head muscles to polyQ aggregation (measured as a ratio of head to body aggregation) in either background. This may potentially indicate that more than one gene in the interval was responsible for the switch in the head muscle susceptibility, or that it depends on SNPs in non-coding RNAs, intergenic regions, or genes with SNPs that were not selected as potentially functionally significant; alternatively, this failure could be due to an inefficient knock-down. On the other hand, RNAi of several genes modified the second phenotype – the overall aggregation of polyQ40 in the body-wall muscle cells. Decreasing expression of two genes, *Y71G12B.23* and *C53H9.3* caused an increase in the number of aggregates in the Q40Bristol animals, with no change in the *drxIR1*;Q40 animals, while knocking down expression of *atg-5* caused a large decrease in aggregation in the *drxIR1*;Q40 strain, with no effect in the Q40Bristol background (**Fig. 4B**). Because reversal of increased aggregation specifically in *drxIR1*;Q40 animals by RNAi is consistent with our genetic analysis for this phenotype in **Fig. 4A**, which suggested that the causative variant in *drxIR1* background will be hypermorphic, this points to *atg-5* as a candidate gene. In agreement, our genome sequencing of *drxIR1*;Q40 strain uncovered two unique SNPs in the 3’UTR of *atg-5* (**Data File 1)**.

Because a hypermorphic effect can be caused by increased expression of the affected gene or protein, and because the SNPs are localized in the regulatory region of *atg-5*, we first measured *atg-5* transcripts. qPCR data revealed no differences in *atg-5* transcript levels in *drxIR1* or *drxIR1*;Q40 animals compared to their respective Bristol strains (**Fig. 4C**). Thus, we asked whether decreasing the protein expression *via* a targeted deletion of *atg-5* could reverse the increased polyQ aggregation in *drxIR1*;Q40 animals, as expected if the variants were hypermorphic. We used *atg-5(bp484)* allele, which has a mutation in a splice donor site of exon 1 disrupting the protein’s expression or function (63, 64). We found that unlike animals that carried one DR1350-derived and one wild-type (Bristol) copy of the interval (*drxIR1/+*;Q40), which we found previously to still exhibit increased aggregation (**Fig. 4A**), *drxIR1* heterozygous animals carrying the *atg-5* mutation in the wild-type interval (*drxIR1/atg-5*;Q40) completely lost the increased aggregation phenotype (**Fig. 4D**). Together, our data suggest that increased levels or activity of ATG-5 protein cause increased polyglutamine aggregation in the body-wall muscle cells.

### Activation of autophagy has divergent effects on polyQ aggregation in different tissues

ATG-5 is an ortholog of the autophagic budding yeast protein ATG5, and of human ATG5. ATG-5 contributes to the initiation of autophagy by forming a complex with LGG-3/ATG12 and ATG-16/ATG16L1, which is recruited to the membrane of the elongating phagophore (65-67), and is required for the lipidation of LGG-1/LC3. Thus, upregulation or activation of ATG-5 by the hypermorphic allele could cause either overactivation or an imbalance in autophagy. Interestingly, ATG5 in mammalian cells can also contribute to the progression of apoptosis, independent of its role in autophagy (68).

Although we saw no increase in the number of GFP::LGG-1 puncta in the muscle cells of *drxIR1* animals under basal conditions (**Fig. 2A**), we did observe more puncta in the hypodermal cells, where autophagy is often scored (69). Thus, we asked whether autophagy the muscle cells was different in *drxIR1* and wild-type (Bristol) animals under activation conditions. We used an autophagy inducer drug, ABT-737, that acts as a BH3-mimetic, inhibiting the antagonistic effects of Bcl-2 (CED-9 in worms) on Beclin-1 (BEC-1) and thus relieving the inhibition of autophagy (70). Treatment with 10µM of ABT-737 indeed induced GFP::LGG-1 puncta in the muscle cells of the wild type (Bristol) animals (**Fig. 5A**). Surprisingly, animals carrying the *drxIR1* interval exhibited an increase in punctate appearance of GFP::LGG-1 protein in the body-wall muscle cells already in response to the DMSO control. Although not previously reported to activate autophagy, low concentrations of DMSO have been reported to extend the lifespan of *C. elegans* and decrease the paralysis associated with Aβ_1-42_ aggregation, when grown in liquid (71, 72). Importantly, ABT-737 resulted in a larger increase in GFP-positive puncta in *drxIR1*;GFP::LGG-1 animals compared to the Bristol background (**Fig. 5A**). These data suggest that *drxIR1* interval increases accumulation of LGG-1/LC31-positive autophagosome structures in response to an activating treatment.

**Fig 5.**
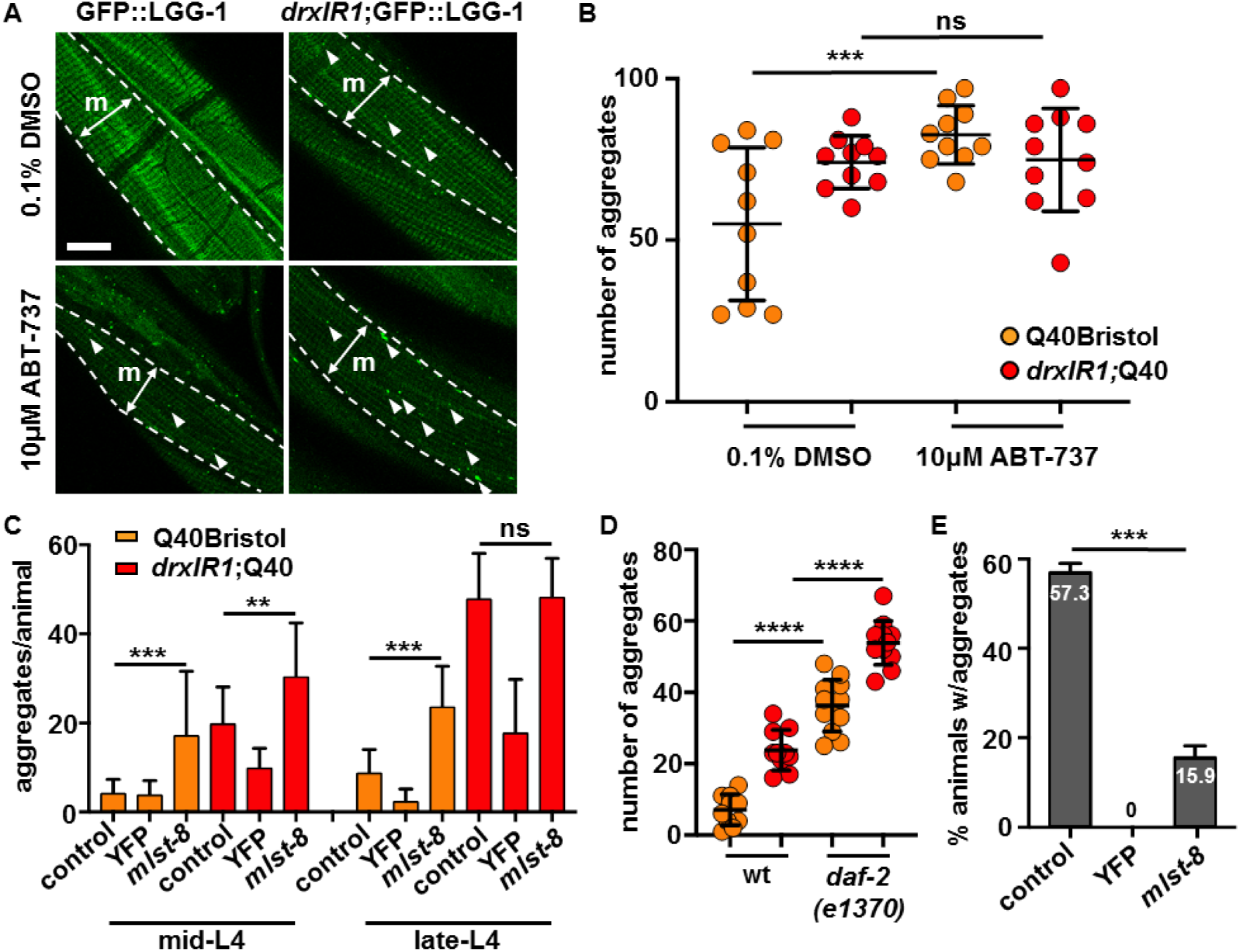
Activation of autophagy has divergent effects on polyQ40 aggregates clearance in different tissues. (**A**) Animals carrying the *drxIR1* interval accumulate more GFP::LGG-1-positive puncta (**arrowheads**) in the body-wall muscle cells upon treatment with autophagy-activating drug ABT-737. Animals were treated with 0.1% DMSO (vehicle control) or 10 µM ABT-737 for 24hr. Shown are confocal projections; one muscle quadrant (**m**) is indicated between punctate lines. Scale bar: 10μm. (**B**) Autophagy-activating drug ABT-737 increases polyQ40 aggregation in the body-wall muscle cells in the wild-type background (Q40Bristol). Aggregation was scored in adult animals, 1 day post L4 (see Methods). Aggregation in the *drxIR1*;Q40 animals is already at maximum under these conditions. Each symbol indicates an individual animal; overlaid are means ± SD. Data were analyzed by ANOVA followed by Bonferroni’s multiple comparisons test, \*\*\**P*=0.0006. (**C**) Activation of autophagy with *mlst-8* RNAi increases aggregation in the body-wall muscles of Q40Bristol mid- or late-L4 animals, and of *drxIR1*;Q40 mid-L4 animals. Aggregation in *drxIR1*;Q40 late-L4 animals is already at maximum. Data are mean ± SD, three independent experiments, 9 to 13 animals per experiment per treatment. Control RNAi was *mec-4*. Data were analyzed by ANOVA followed by Bonferroni’s multiple comparisons test, \*\*\**P*=0.0007, \*\**P*=0.0082. (**D**) Introduction of the *daf-2(e1370)* allele increases polyQ40 aggregation in the body-wall muscles in both Q40Bristol and *drxIR1*;Q40 animals. Aggregation was scored at mid-L4. Each symbol indicates an individual animal; overlaid are means ± SD. Data were analyzed by ANOVA with Bonferroni’s multiple comparisons test, \*\*\*\**P*<0.0001. (**E**) Activation of autophagy with *mlst-8* RNAi strongly suppresses polyQ aggregation in the intestinal cells. Percent of animals with Q44::YFP aggregates in the intestine of day 4 adult were scored, as in refs(73, 74), for each indicated RNAi treatment. Control RNAi was *mec-4.* Data are mean ± SD. Data were analyzed by ANOVA followed by Bonferroni’s multiple comparisons test, \*\*\**P*=0.0003.

The larger increase in LGG-1 puncta in *drxIR1*;GFP::LGG-1 animals could indicate that *atg-5* hypermorphic allele causes either a stronger activation of autophagy, or slower lysosomal degradation. Because autophagy is known to promote clearance of polyglutamine aggregates (75), the increased aggregation in *drxIR1* background appeared consistent with slower degradation, while activation of autophagy would have been expected to decrease aggregation (76). To confirm this, we asked whether activation of autophagy with ABT-737 indeed decreased polyQ aggregation in the wild type (Bristol) background. Surprisingly, treatment of Q40Bristol animals with this autophagy activator resulted in a large increase, rather than decrease, of polyQ40 aggregation in the body-wall muscles, with ABT-737-treated animals exhibiting a 44% increase in the number of aggregates (**Fig. 5B**). These data suggest that, counter to expectations, activation of autophagy may enhance polyglutamine aggregation. We did not detect a further increase in aggregation in *drxIR1* background, since the drug treatment protocol dictated scoring aggregates in young adult animals (see materials and methods), when aggregation in *drxIR1*;Q40 is already close to maximal.

Because this effect of autophagy was unexpected, and because the drug treatment may not be reliable in *C. elegans*, we tested two different genetic approaches known to activate autophagy to confirm these findings. Each of the two approaches activates autophagy *via* mechanisms distinct from that of ABT-737. First common approach is inactivation of mTOR (77). In *C. elegans*, inactivation of LET-363/mTOR indeed activates autophagy, as shown by increase in GFP::LGG-1 puncta (78). However, inactivation of LET-363 also causes larval arrest (79), which itself will affect polyQ aggregation. To overcome this, we targeted mTOR interacting protein MLST-8/mLST8, which is required for the kinase activity of mTOR (80), but can be downregulated in *C. elegans* without causing larval arrest (81). RNAi knock-down of *mlst-8* resulted in a 1.6-fold increase in polyQ40 aggregation in Q40Bristol animals (**Fig. 5C**, late-L4). Similar to the results of the drug treatment, *mlst-8* RNAi had no significant effect in *drxIR1*;Q40 animals. We asked whether the apparent lack of effect on the *drxIR1*;Q40 animals was indeed due to the already high aggregate numbers at this developmental stage, by repeating the RNAi in younger animals, and observed an even stronger, 3-fold, increase in polyQ40 aggregation in Q40Bristol animals, and a 1.5-fold increase in *drxIR1*;Q40 animals (**Fig. 5C**, mid-L4).

As a second genetic approach, we tested the effect of decreased activity of insulin/IGF-like signaling pathway, since reduction of function of the sole *C. elegans* orthologue of insulin/IGF receptor, DAF-2, is known to cause activation of autophagy, including in the body-wall muscle cells (52, 82). Introduction of the hypomorphic *daf-2(e1370)* allele caused a 5.1-fold increase in aggregates in the Q40Bristol background, and 2.3-fold further increase in *drxIR1*;Q40 animals (**Fig. 5D).** The increase in polyQ aggregation in *daf-2(e1370)* background is consistent with previous reports (83). Together, these pharmacological, RNAi, and genetic data suggest that aggregation of polyQ40 in the body-wall muscle cells is paradoxically increased by activation of autophagy.

Previous studies indicate that autophagy levels, both basally and in response to a trigger, can be different in different *C. elegans* and mammalian tissues (51, 84). Intriguingly, in these reports, mouse slow-twitching muscles showed nearly no basal autophagy and only low to moderate induction in response to starvation (84), and *C. elegans* body-wall muscle cells also showed low basal autophagy compared to other tissues (51, 52), consistent with our observation of the low numbers of LGG-1-positive puncta in the muscle cells in **Fig. 2C**. Thus, we asked whether activation of autophagy may have a different effect on polyQ aggregation in muscles than in a different tissue. In addition to the muscle-expressed polyQs, the neuronal and intestinal fluorescent polyQ models have been described in *C. elegans* (73, 85). Since aggregation of the moderate polyQ40 expansions in neurons is only detectable by FRAP, we chose to examine the intestinal polyQ44 model. We found that unlike in the muscle-expressed polyQ model, activation of autophagy *via* RNAi knock-down of *mlst-8* resulted in a large (3.5-fold) decrease in the percentage of animals exhibiting polyglutamine aggregation in intestine (**Fig. 5E**). Thus, depending on the tissue, activation of autophagy can either clear polyglutamine aggregates or cause their accumulation.

## Discussion

Using natural genetic variation, we identified an unexpected divergence in how activation of autophagy in different tissues impacts the behavior of aggregation-prone polyglutamine expansions. It is broadly appreciated that autophagy can be both protective and detrimental to cells and organisms (86). For example, ER stress-induced autophagy is protective in cancer cells but contributes to apoptosis in non-transformed cells (87), while starvation-triggered autophagy in *C. elegans* pharyngeal muscle can switch from protective to pro-death, depending on its level of activation (50). However, with respect to the clearance of misfolded aggregated proteins, activation of autophagy is generally considered to be a positive, protective response (88, 89). Therefore, activation of autophagy has been thought of as a nearly universal therapeutic approach to neurodegenerative diseases caused by protein aggregation (90). While we did not test the neuronal polyQ models, the divergence in how polyQ expansions in intestinal and muscle cells respond to activation of autophagy suggests that interplay between autophagy and protein aggregation depends on the cellular context. We find that both the natural variants in *atg-5*, and the more traditional genetic and pharmacological ways of activating autophagy, increased rather than decreased polyQ aggregation in the muscle cells of *C. elegans*. This represents a striking departure from the current paradigm. On the other hand, polyQ aggregation in the intestinal cells, as expected, was decreased by the same treatment. Considering the significant involvement of skeletal muscle and non-neuronal tissues in HD and other polyglutamine diseases, including the induction of the muscle catabolic phenotype and muscle wasting (91-95), a more nuanced understanding of integration of autophagy with cellular physiology is needed.

The use of natural variation was instrumental in uncovering this unexpected cell-specific effect of autophagy on protein aggregation. The DR1350-derived variants that we identified as being responsible for the increased aggregation of polyQ40 in the muscle cells are in the regulatory 3’UTR region of the *atg-5* gene. Our genetic analysis points to the gain of expression as the mechanism of *atg-5* variants. Based on the ability of one additional copy of the wild-type, Bristol-derived *atg-5* to mimic the effect of these natural variants (**Fig. 4A**), and because deletion of one copy of *atg-5* reverses the effect of the variants in the remaining copy (**Fig. 4D**), we estimate that the variants increase the expression of ATG-5 protein by less than 2-fold. Strikingly, introduction of one additional Bristol-derived copy of *atg-5* into the animals already carrying two DR1350-derived hypermorphic alleles, increases the polyQ aggregation even further, to about 6-fold above normal. This indicates a quantitative relationship between the levels of ATG-5 protein and increased polyQ aggregation in the muscle. Although we are currently unable to directly modulate autophagy in *C. elegans* in a graded manner, the ability of three distinct methods of activating autophagy to mimic the effect of the variants argues that the increase in ATG-5 affects the polyQ aggregation by increasing autophagy, rather than for example by causing stoichiometric imbalance and autophagy inhibition (76), or coupling to apoptosis pathway (68). The precise mechanistic basis of this quantitative relationship will need to be investigated further.

One important aspect of our findings is the cryptic nature of the modifier variants in *atg-5*. Cryptic variation typically does not cause phenotypic changes on its own, but becomes phenotypically “exposed” when challenged with a stressful environment, thus contributing to disease susceptibility (96-98). Polyglutamine expansions may mimic cellular stress, for example by destabilizing the folding environment (37) or disrupting transcriptional control (99). Indeed, the *atg-5* variants identified here as modifiers are derived from a phenotypically normal wild strain DR1350, and we did not detect significant alterations in the basal autophagy in the muscles of *drxIR1* animals unless they were challenged with the aggregation-prone polyQ40, or with autophagy-activating drug ABT-737.

In addition to being exposed by stress, the phenotypic expression of cryptic modifier variants may reflect their more direct interactions with the disease-causing mutation. For example, in humans, analysis of HD modifier loci on Chromosomes 8 and 15 showed that these variants influence certain clinical readouts in subjects with expanded polyQ tracts, prior to the appearance of disease symptoms, while they have no major effects in control individuals without expansions (18). The suspected culprit for the modifying effect of the Chromosome 15 locus, the DNA endo/exonuclease FAN1, may be changing the disease phenotypes or age of onset by directly affecting the stability of the polyQ-encoding repeat in somatic tissues (18, 100).

Interestingly, the HD disease progression study (18) suggested that the modifiers could have distinct modifying effects in different cell populations. In our study, the cryptic nature of the *atg-5* variants allowed detection of the unusual cell-specificity in autophagy – aggregation relationship, because non-cryptic genetic variants that ectopically activate autophagy already under basal conditions have additional strong phenotypes that can mask the effects of activated autophagy on polyQ aggregation. For example, loss of function of *C. elegans* mTOR leads to larval arrest (79); hypomorphic mutations in insulin/IGF signaling pathway, in addition to activating autophagy, trigger numerous other developmental, stress responsive, and metabolic pathways (101-103) that can have their own effects on the aggregation-prone protein; and even non-genetic means such as activation of autophagy by nutrient deprivation are accompanied by the metabolic and protein expression changes (104) that can mask the more specific effect on the polyQ behavior. Natural variation may thus indeed identify the candidate modifier pathways and mechanistic relationships in aggregation diseases that are distinct from those identified by the traditional approaches.

The reasons the muscle cells are differentially sensitive to autophagy with respect to protein aggregation, or why this is not true for other aggregation-prone proteins, are not yet known. The selectivity of aggregation effects of autophagy towards the polyglutamine expansions would argue against a global dysregulation of protein homeostasis in the muscle cells of *drxIR1*animals, which is supported by our data. It is possible however, that ectopic activation of autophagy disrupts select proteostasis processes that impinge on the polyQ aggregation or clearance in these cells. Another possibility is that autophagic degradation of polyQ expansions requires a specific “signal” or adaptor, which may be competed away during generic increase in autophagy in the muscle cells, but remains sufficient in the intestine. The polyQ-expanded huntingtin protein (Htt) indeed requires specific adaptors, such as Tollip, to be cleared by autophagy (105), although whether this is also true for polyQ expansions outside the Htt context is not clear. Yet another possibility is that polyQ expansions themselves interfere with autophagy. For example, polyQ-expanded Htt have been suggested to interfere with the delivery of cargoes to autophagic vacuoles (106), and shown to co-aggregate with the autophagy adaptor Tollip, potentially disrupting other functions of this multi-tasking protein (105). If so, the low basal levels of autophagy may render the proteostasis of the muscle cells to be more sensitive to the polyQ expansions.

Muscle cells may also have a different regulation of or dependence on autophagy because autophagy of the muscle is an adaptive response of many metazoans to starvation (107). While basal autophagy is important for muscle maintenance, not only deficiency in autophagy but also its over-activation can lead to muscle atrophy (108-110). Indeed, in *C. elegans*, body-wall muscles in young animals have low basal levels of autophagy relative to other tissues (51, 52), while in mice, the slow-twitching (soleus) muscles exhibited little induction of autophagy after 24 hours of starvation, as defined by the autophagosome counts, distinct from the fast-twitching (extensor digitorum longus) muscles that had significant induction (84). Moreover, the distribution of autophagosomes was different between the fast- and slow-twitching muscle types, supporting the idea of differential autophagy regulation in different cells or tissues.

In addition to the traditional mouse models, the genetic model systems such as worm, fly and yeast, in which natural variation can be readily combined with modeling the gain-of-function mutations by transgenesis, offer new opportunities to identify the cryptic modifier pathways for neurodegeneration and protein misfolding and aggregation (10, 111-115). Examples of this approach include a recent study in *C. elegans* that showed that the ability of α-synuclein to cause transcriptional and phenotypic changes is substantially modified by the genetic background (40), and a study using a Drosophila Genetic Reference Panel (116), that uncovered an unexpected role of heparin sulfate protein modifications in modifying the toxic effects of the misfolded mutant of human insulin, a cause of permanent neonatal diabetes (117). The important feature of the cryptic modifier pathways that can be identified by these approaches is that they harbor natural variants shaped by selection, and thus will pinpoint the naturally plastic potential genes and networks (14), amenable to pharmacological manipulation without negative effects on the organism.

## Materials and Methods

### Nematode strains and growth conditions

Nematodes were grown at 20°C on nematode growth medium (NGM) plates, seeded with *E. coli* OP50 (118). Animals were synchronized by picking gastrula stage embryos onto fresh plates, unless otherwise noted.

The following stains were obtained from *Caenorhabditis* Genetics Center (CGC): AM141 [*rmIs333*(p*unc-54*::Q40::YFP)], AM140 [*rmIs132*(p*unc-54*::Q35::YFP) I], CF1038 [*daf- 16(mu86)* I], TJ356 [*zIs356(*p*daf-16::daf-16a/b*::GFP;*pRF4(rol-6(su1006)*) IV], PS3551 [*hsf-1(sy441)* I], DA2123 [*adIs2122*(p*lgg-1*::GFP::*lgg-1* + *rol-6(su1006)*)], KR1108 [*unc-11(e47) dpy-5(e61)* I], KR292 [*him-1(h55) dpy-5(e61) unc-13(e450)* I; sDp2 (I;f)], MT1434 [*egl-30(n686)* I], and CB1370 [*daf-2(e1370)* III]. TGF205 [*xzEx3*(p*unc-54*::UbG76V::Dendra2)] was made by crossing out *glp-1(e2141)* from AGD1033.

The AS408 [p*unc-54*::GFP::UNC-54], AM583 [*rmIs249*(p*let-858::hsf-1;*p*myo-2::*GFP)], AM738 [*rmIs297*(p*vha-6*::Q44::YFP;*rol-6(su1006)*)], and AM930 [*rmIs335*(p*unc-54*::Aβ(1-40)::CFP)] strains were kindly provided by the Morimoto lab, and the HZ1732 [*atg-5(bp484)* I*;him-5*] strain by the Colón-Ramos lab. The Q40DR1350 and recombinant inbred lines (RILs) 2, 12, 12(2) and 15 were described in (38).

The *drxIR1*(I, DR1350>Bristol) locus and/or the Q40 locus were introduced by genetic crosses into the following strains: TGF134 [*drxIR1*;*rmIs333*(p*unc-54*::Q40::YFP)], TGF130 [*drxIR1*;p*unc-54*::GFP::UNC-54], TGF353 [*drxIR1*;*adIs2122*(p*lgg-1*::GFP::*lgg-1* + *rol-6(su1006)*)], TGF208 [*xzEx3*(p*unc-54*::UbG76V::Dendra2);*rmIs333*(p*unc-54*::Q40::YFP)], TGF207 [*drxIR1*;*xzEx3*(p*unc-54*::UbG76V::Dendra2);*rmIs333*(p*unc-54*::Q40::YFP)], TGF088 [*daf-16(mu86)* I; *rmIs333*(p*unc-54*::Q40::YFP)], TGF188 [*drxIR1*;*daf-16(mu86)* I;*rmIs333*(p*unc-54*::Q40::YFP)], TGF086 [*zIs356(*p*daf-16::daf-16a/b*::GFP;*pRF4(rol-6(su1006)*) IV;*rmIs333*(p*unc-54*::Q40::YFP)], TGF190 [*drxIR1*;*zIs356(*p*daf-16::daf-16a/b*::GFP;*pRF4(rol-6(su1006)*) IV;*rmIs333*(p*unc-54*::Q40::YFP)], TGF187 [*hsf-1(sy441)* I;*rmIs333*(p*unc-54*::Q40::YFP)], TGF170 [*drxIR1*;*hsf-1(sy441)* I;*rmIs333*(p*unc-54*::Q40::YFP)], TGF036 [*rmIs249*(p*let-858::hsf-1;*p*myo-2::*GFP);*rmIs333*(p*unc-54*::Q40::YFP)], TGF189 [*drxIR1*; *rmIs249*(p*let-858::hsf-1;*p*myo-2::*GFP);*rmIs333*(p*unc-54*::Q40::YFP)], TGF203 [*drxIR1*;*rmIs335*(p*unc-54*::Aβ(1-40)::CFP)], TGF342 [*drxIR1*;*rmIs132*(p*unc-54*::Q35::YFP) I], TGF261 [*rmIs333*(p*unc-54*::Q40::YFP);*him-1(h55) dpy-5(e61) unc-13(e450)* I; sDp2 (I;f)]TGF275 [*drxIR1*;*rmIs333*(p*unc-54*::Q40::YFP);*him-1(h55) dpy-5(e61) unc-13(e450)* I; sDp2 (I;f)], TGF089 [*daf-2(e1370)* III;*rmIs333*(p*unc-54*::Q40::YFP)]; TGF127 [*drxIR1*;*daf-2(e1370)* III;*rmIs333*(p*unc-54*::Q40::YFP)].

The *drxIR1*;Q40 strain was made by the following scheme: Q40Bristol males were mated to RIL2 hermaphrodites, and 5-10 F1 hermaphrodite progeny, identified by the lack of RIL2-like increased head aggregation phenotype, were picked onto fresh plates. F2 generation was examined for the expected 1:3 segregation of the increased head aggregation phenotype, and 7-10 F2 hermaphrodites with this phenotype were further mated with Q40Bristol males. This mating-selection cycle was repeated 23 times. The resulting strain was named *drxIR1;*Q40.

The introduction of *drxIR1* locus by genetic crosses was confirmed by detecting the presence of the SNP 5 (WBVar00016276) (**Suppl. Fig. 1**): a 743bp fragment containing the variant was amplified using the *drxIR1* primers (**Suppl. Table 3**), at an annealing temperature of 60°C, to produce an amplicon of 743bp, and the PCR product was digested with SalI. The SalI site is present in the Bristol background, producing 432bp and 311bp products after the digest, but is absent in the DR1350 background.

### Genome sequencing

Genomic DNA from *drxIR1*;Q40 and Q40Bristol was extracted from flash frozen pellets of a mixed populations, using phenol:chloroform (Sigma, USA). DNA was sequenced using the NextSeq 500 System (Illumina, USA) at the Wistar Institute (Philadelphia PA, USA). Sequencing data was analyzed using the Galaxy (119) CloudMap pipeline as described in (42), and WS220 genome assembly. The CloudMap SnpEff tool was utilized to annotate the genetic variants and predict their functional effects on genes and proteins (62). SNPs with the following annotations were considered as potentially functionally-significant: non-synonymous coding, start gained or lost, stop gained or lost, splice site donor/acceptor, frameshift, and 5’ or 3’ UTR.

### Quantification of polyQ40 aggregation

Aggregation was scored by counting fluorescent foci in images collected from animals immobilized with 20 mM NaN_3_, using a Leica M205FA stereoscope with a digital camera (Hamamatsu Orca R2). For synchronization, 15-20 well-fed L4 animals from non-crowded plates were transferred to new plates, gastrula stage embryos were picked 2-3 days later, and hatched animals were allowed to develop for specified time or to specified developmental stage. Aggregation was scored in late-L4 animals, unless otherwise indicated. The developmental larval stage was confirmed based on the germline development, or by days since L4 (for older adults). For data expressed as means, the number of animals for each data point is indicated in the figure legends.

### Microscopy

For confocal images, animals were immobilized on 2% agar pads with 20 mM NaN_3_ and imaged with Zeiss LSM700 microscope at Cell Imaging Center, Drexel University. Z-stacks were acquired at 0.4 µm intervals as 12-bit images, using 63x 1.4NA objective, and analyzed with ImageJ. For the quantification of autophagic vesicles, Z-stacks were collapsed as maximum intensity projections, the muscle cells were outlined, and the GFP::LGG-1-positive puncta within the outlined cells were counted. 30 to 40 cells from 8 to 10 L4 animals were analyzed per genotype. To compare GFP::UNC-54 protein levels, GFP fluorescence was measured within the same size area (∼9μm^2^) in the center of each analyzed muscle cell, over the myofilaments. 16-20 cells from 4-5 animals per genotype were measured. An identical size area measured away from the myofilaments was used for background subtraction.

Fluorescent recovery after photobleaching (FRAP) was performed on day 2 adults (for aggregated Q40) and L4 larvae (for soluble Q40) animals, as in (85), using the Zeiss LSM700 confocal microscope. Photobleaching was performed with 488 nm laser, by 100 iterations at 100% laser power. Imaging during recovery was at 0.2% power. Relative fluorescence intensity (RFI) was determined with the following equation: RFI = *(T*_*t*_*/C*_*t*_*)/(T*_*0*_*/C*_*0*_), with *T*_*0*_ representing the total intensity of the region of interest before photobleaching and *T*_*t*_ the intensity in the same are at any time after. We normalized against an unbleached area in the same cell, where *C*_*0*_ is a control area before bleaching and *C*_*t*_ represents any time after bleaching (85). 7-18 aggregates from 3 animals each were measured per strain for aggregated Q40, and 5 cells from 2 animals each were measured per strain for the soluble Q40 controls.

For stereo images, animals were immobilized on NGM plates in a drop of 20 mM NaN_3_. Imaging was performed using a Leica M205FA stereo microscope with an Orca R2 digital camera (Hamamatsu). The magnification and the intensity of fluorescent sources (Chroma PhotoFluor 2) were kept constant within experiments. UbG76V::Dendra2 animals were imaged with a narrow-bandpass CFP filter (Chroma), to avoid the spectral overlap with the Q40::YFP protein.

### Native protein extracts

To prepare native protein extracts, synchronized embryos were prepared by hypochlorite treatments and larvae were collected once they reached the L3 stage. The worm pellets were mechanically disrupted and lysed in 0.5% Triton-X 100 buffer as described in (39). For SDS solubility, native protein extracts were incubated in 5% SDS for 15 min at room temperature prior to running on a 5% continuous native-PAGE, at 25 mg of total protein per lane. Gels were imaged on a Typhoon FLA7000 scanner (General Electric, USA) with ImageQuant TL software to quantify YFP fluorescence. All experiments were performed three times.

### qPCR

∼50μl pellets of L4 stage worms were flash frozen in liquid nitrogen and RNA extraction was performed using TRIzol (Life Technologies, USA) and chloroform (Sigma, USA) reagents. The samples were treated with DNase (DNA-*free*, Life Technologies, USA) to remove any genomic DNA, and iScript cDNA synthesis kit (Bio-Rad, USA) was used to reverse transcribe 1-2 µg of RNA per sample. The expression of selected genes was measured using iTaq Universal SYBR Green Supermix (company) and the ViiA detector (Applied Biosystems). Each biological replicate was run in triplicate, and data analyzed using the δδCT method (120). Three biological replicates were used to assess statistical significance. Tubulin (*tbg-1*) was used as the internal control, as it was stable between the *drxIR1* and the Bristol strains. *tbg-1* primers were obtained from (121). Primer sequences are listed in **Suppl. Table 3**.

### RNAi experiments and constructs

For RNAi experiments, NGM plates containing 100 µg/ml ampicillin and 0.4 mM IPTG were seeded with control (L4440 empty vector, unless otherwise noted) or experimental overnight RNAi bacterial cultures and incubated at room temperature for 2 days prior to plating worms. Nematodes were cultured on the RNAi plates from gastrula stage embryos for two generations. RNAi strains were from the Ahringer library (J. Ahringer, University of Cambridge, Cambridge, U.K.), except for those corresponding to *mab-20, Y71G12B.18, Y71G12B.33, Y71G12B.23, Y71G12B.35, drag-1, Y71G12B.31, ubc-3, tln-1, Y71G12B.25, pghm-1, C53H9.3, tag-96, tub-2, Y51F10.4, and spe-48*; these were made by cloning a unique 0.8 to 1.2 Kb fragment from each gene into the L4440 plasmid and transforming into the *E. coli* strain HT115. Primer sequences are listed in **Suppl. Table 4**. All experiments were repeated three times, the total (combined) number of animals is indicated in figure legends.

### ABT-737 treatment

20-40 gastrula stage embryos were grown on OP50 bacteria for two days at 20°C, nematodes collected, washed, and exposed to either 0.1% DMSO (Sigma, USA) as solvent control, or 10 µM ABT-737 (ApexBio, Taiwan). Earlier exposure to ABT-737 resulted in larval arrest. Animals were incubated in the drug solution with shaking for 24 hours, pipetted onto plates, and either scored for aggregation or imaged.

### Statistical analyses

ANOVA and *t*-test analyses were performed with Prism software (GraphPad, USA), using α value of 0.5. ANOVA was followed by Bonferroni’s multiple comparisons post-test. All p-values and significance levels are indicated in the figures and figure legends.

## Acknowledgements

We thank the Colón-Ramos lab at Yale School of Medicine and the Morimoto lab at Northwestern University for contributing worm strains. Some strains were provided by the *Caenorhabditis* Genetics Center (CGC) at the University of Minnesota, which is funded by NIH Office of Research Infrastructure Programs (P40OD010440). We would like to thank Anna Lysenko for experimental assistance.

## Supplemental Files

**Suppl. Fig 1.**
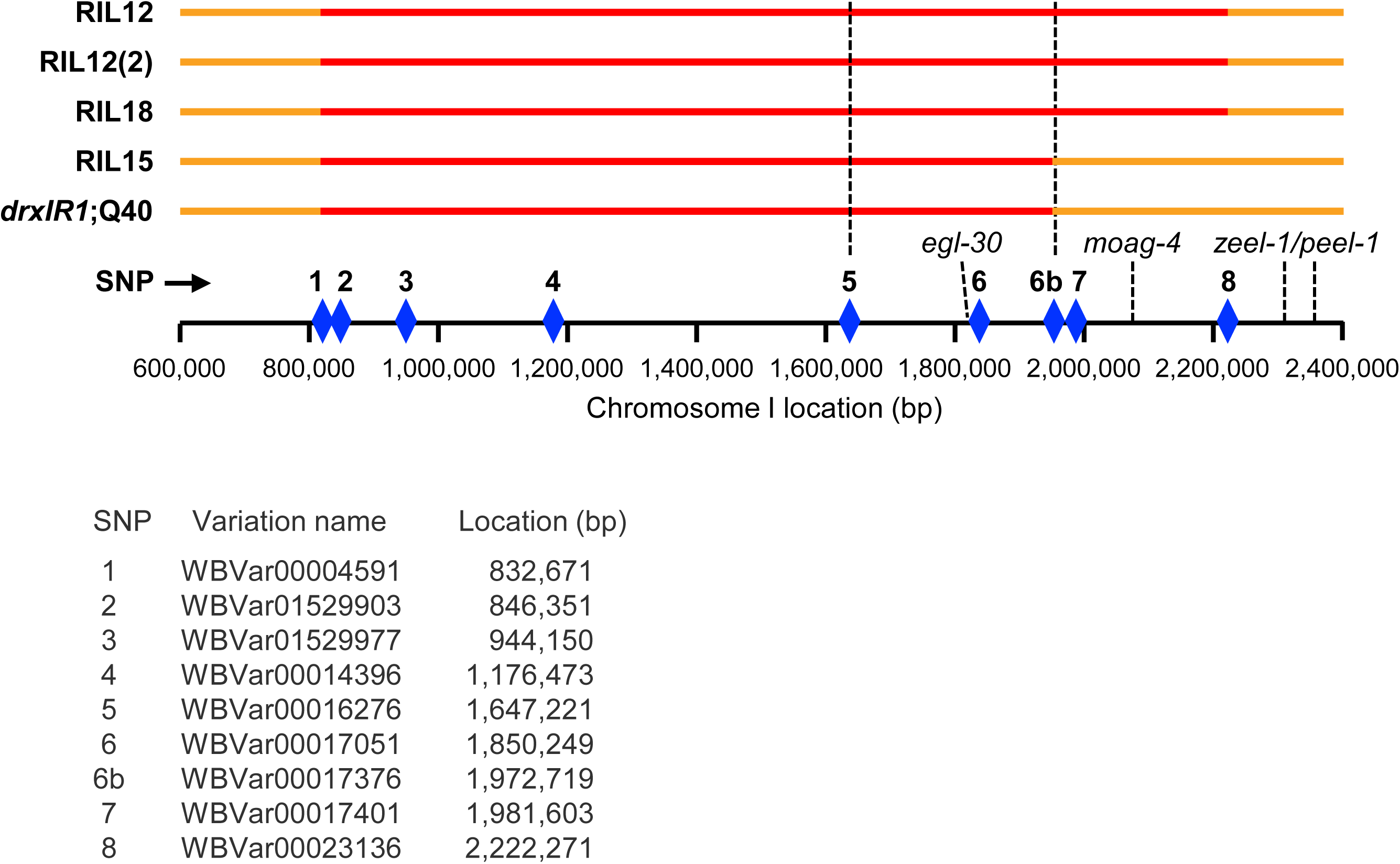
Schematic of the *drxlR1*interval and SNPs used for mapping. **Red**: the 1.4Mb genomic region on Chromosome I containing the DR1350-derived intervals in the RIL2-derived *drxlR1*;Q40 strain and the four remaining high aggregation RILs (RIL12, RIL12(2), RIL18 and RIL15); **orange**: the Bristol background. **Punctate lines** delineate the narrowed 326 Kb interval containing the candidate genes tested by RNAi. **Diamonds**: SNPs used to confirm the presence of the interval; SNP 6b (ChrI:1,972,719 (WBVar00017376)) is Bristol-derived in *drxIR1*;Q40 and RIL15 animals. **L**ocations of *egl-30, moag-4* and the incompatibility locus *zeel-1/peel-1*are also indicated. The coordinates here correspond to the WormBase release WS270 (122).

**Suppl. Fig 2.**
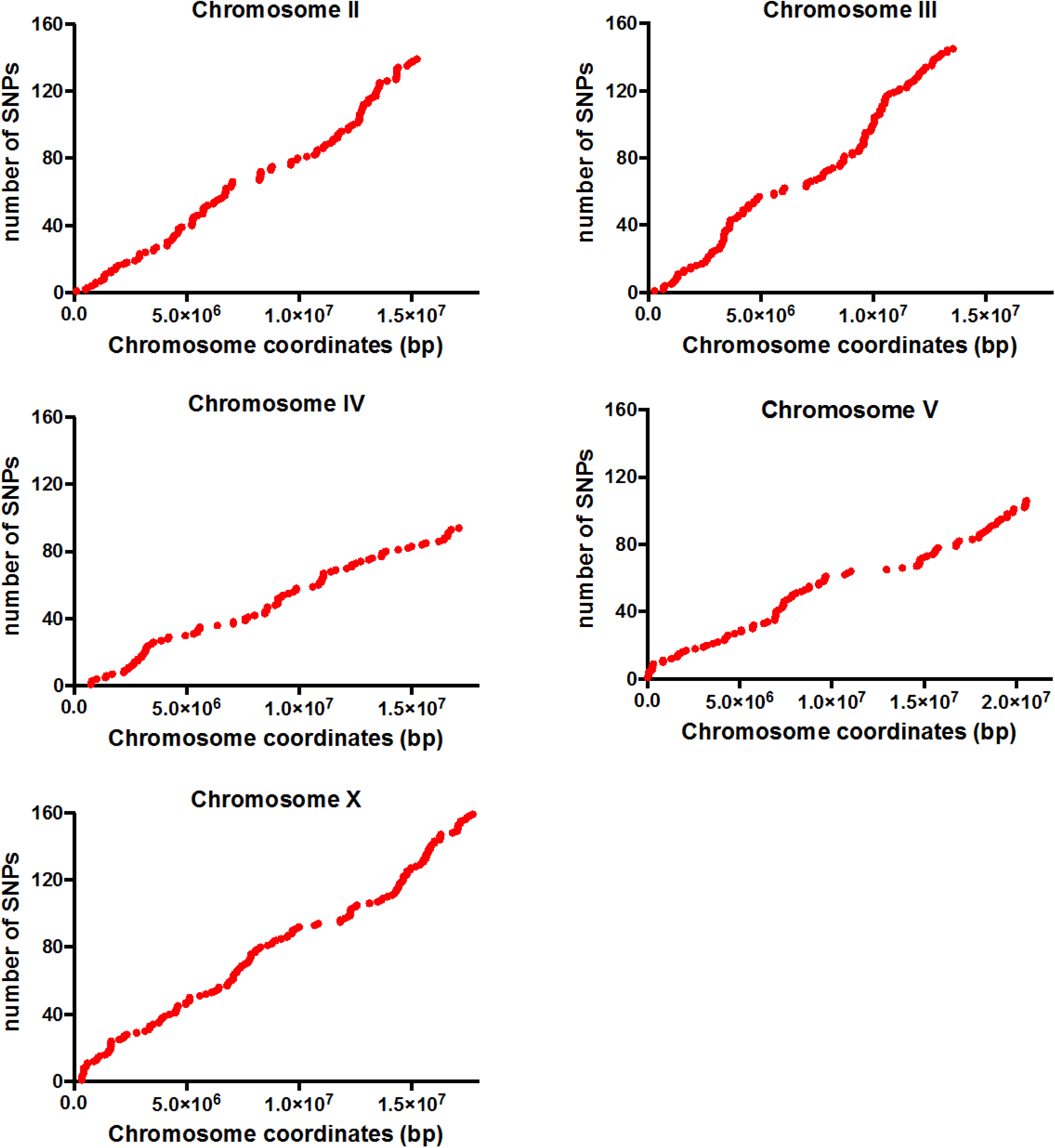
Cumulative distribution of unique SNPs across remaining Chromosomes. Chromosomes II through X in the *drxlR1*;Q40 strain accumulated up to 160 unique SNPs each. Shown are SNPs remaining after subtraction of the variants present in the parental Q40Bristol strain and the known Hawaii SNPs.

**Suppl. Fig 3.**
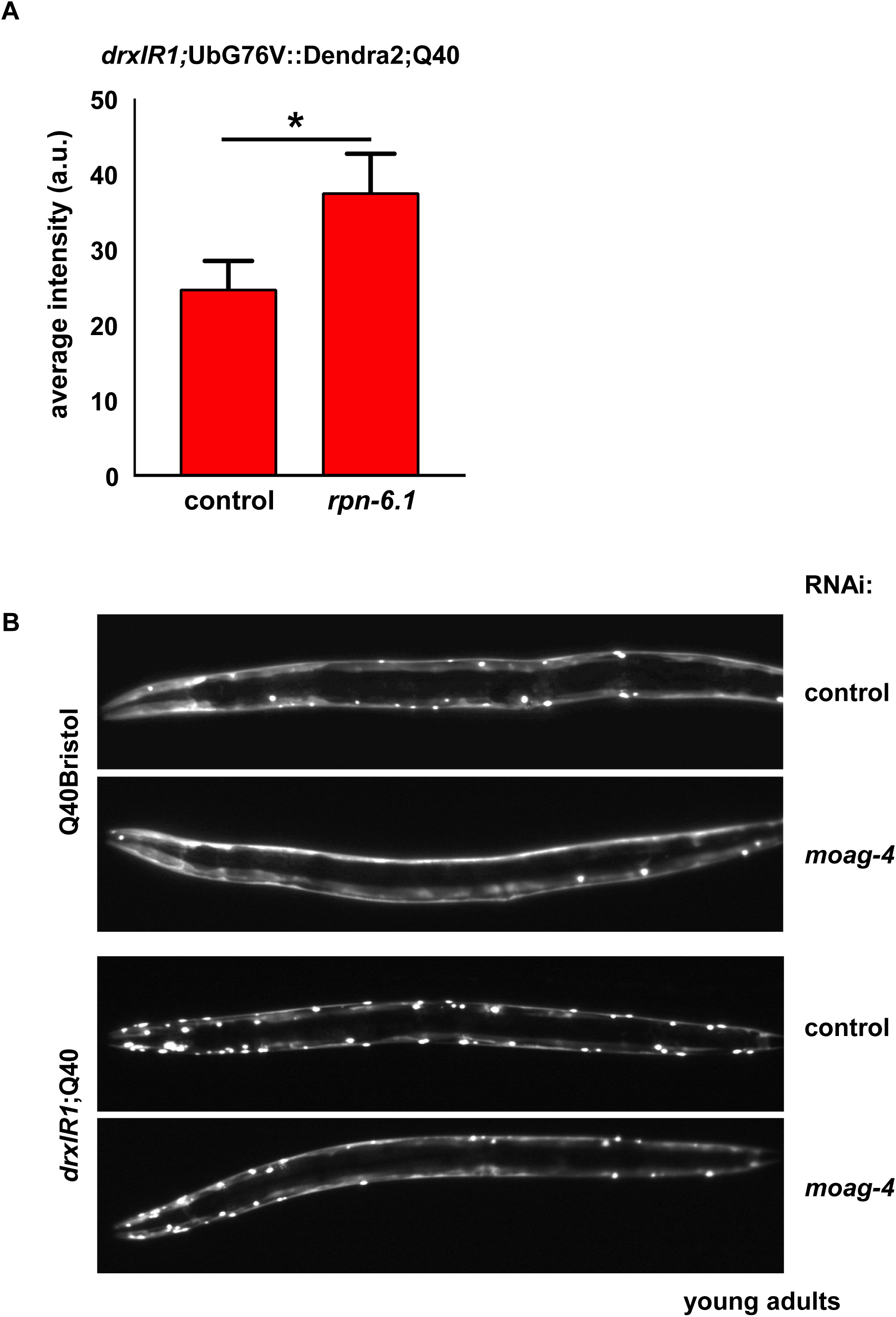
Controls for basal proteostasis effects of the *drxIR1* locus. (**A**) The UbG76V::Dendra2 proteasome reporter is sensitive to decreased proteasome levels. Knockdown of a proteasome subunit *rpn-6.1*, via RNAi, increased the average intensity of the Dendra2 compared to control treatment. Images were taken and quantified as in Fig. 2D. Data are mean ± SD. Data were analyzed by unpaired *t*-test, two-tailed, \**P*=0.0244. (**B**) Stereomicrographs of **young adult** animals after treatment with control or *moag-4* RNAi. *moag-4* RNAi decreased aggregation in both backgrounds, but preserved the increased aggregation *drxIR1*;Q40 animals relative to Q40Bristol.

**Suppl. Table 1.** Loss-of-function analysis for the RIL2-like head aggregation phenotype. sDP2 free duplication covers most of the left arm of Chromosome I, extending through *dpy-5* marker but not through *unc-13. drxIR1*;Q40 animals were crossed with KR292 [*him-1(h55);dpy-5(e61);unc-13(e450)*I; sDp2(I;f)], F1 progeny that either did (based on segregation of *unc* non-*dpy* phenotype among their progeny) or did not inherit the sDp2 duplication were singled, and their F2 progeny scored for the increased ratio of head to body aggregation (RIL2-like) and the dumpy phenotypes. The RIL2-like phenotype behaved genetically as did the known loss-of-function *dpy-5(e61)* allele.

| # of F1 animals | # of F2 animals scored | # RIL2-like F2 animals | # *dpy* F2 animals | % RIL2-like F2 animals | % *dpy* F2 animals |
| --- | --- | --- | --- | --- | --- |
| 22 | 4533 | 1041 | 1109 | 23 | 24.5 |

**F1 animals that *did* inherit sDp2**
| # of F1 animals | # of F2 animals scored | # of RIL2-like F2 animals | # of *dpy* F2 animals | %RIL2-like F2 animals | % *dpy* F2 animals |
| --- | --- | --- | --- | --- | --- |
| 5 | 707 | 91 | 91 | 12.8 | 12.8 |

**Suppl. Table 2.**
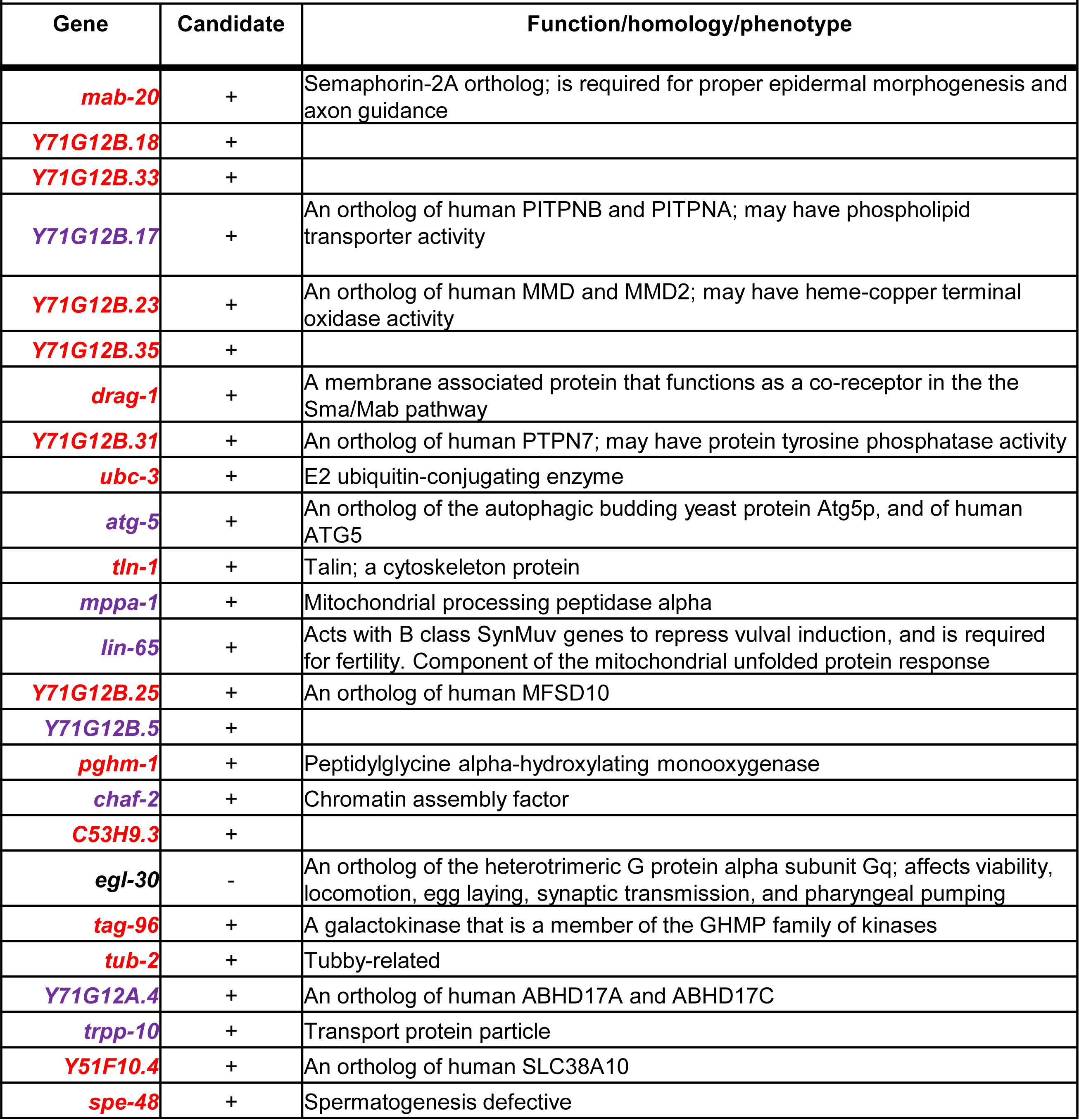
Candidate genes tested by RNAi. 24 candidate genes present in the target 326 Kb of *drxIR1* interval (between SNPs 5 and 6b (**Suppl. Fig. 1**)) are indicated in color. Genes were defined as candidates based on the SnpEff annotations (see Methods and **Data File 1**). *egl-30* was excluded based on genetic crosses. Genes in **purple** were targeted by clones from the Ahringer RNAi library. RNAi targeting constructs for genes in **red** were prepared in this work.

**Suppl. Table 3.**
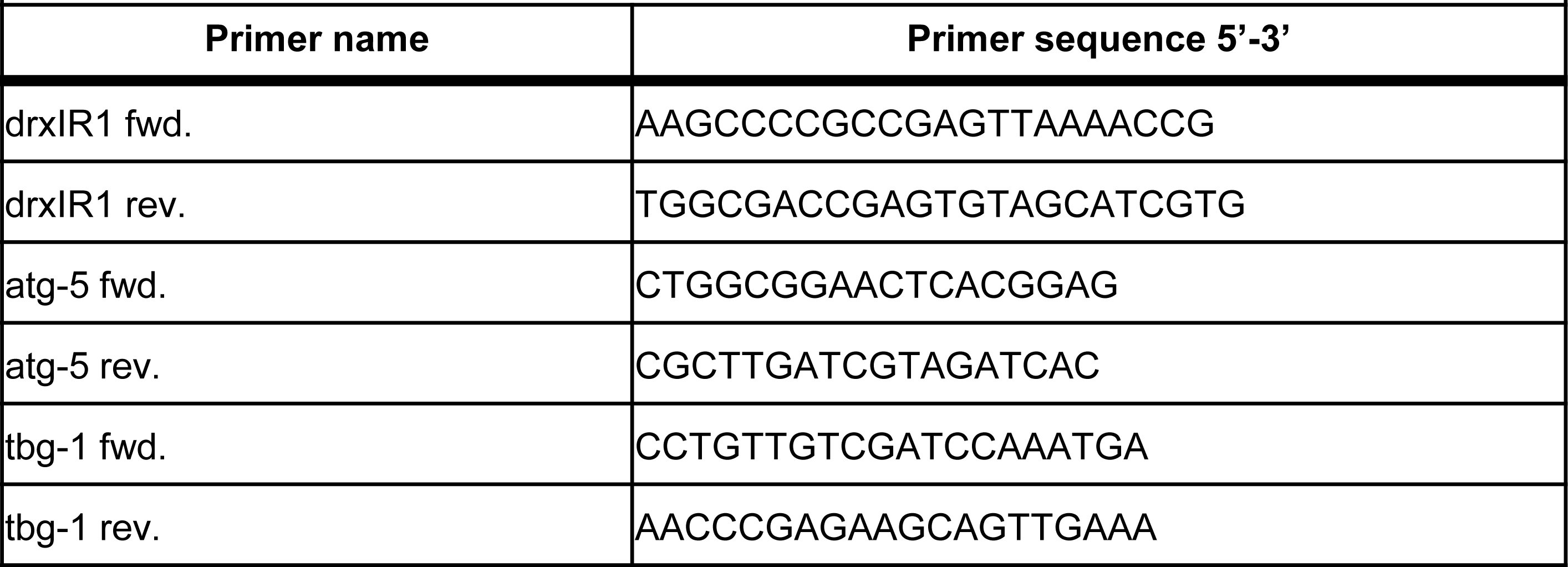
Primers used for genotyping the *drxIR1* locus and for qPCR analysis of *atg-5* expression.

**Suppl. Table 4.**
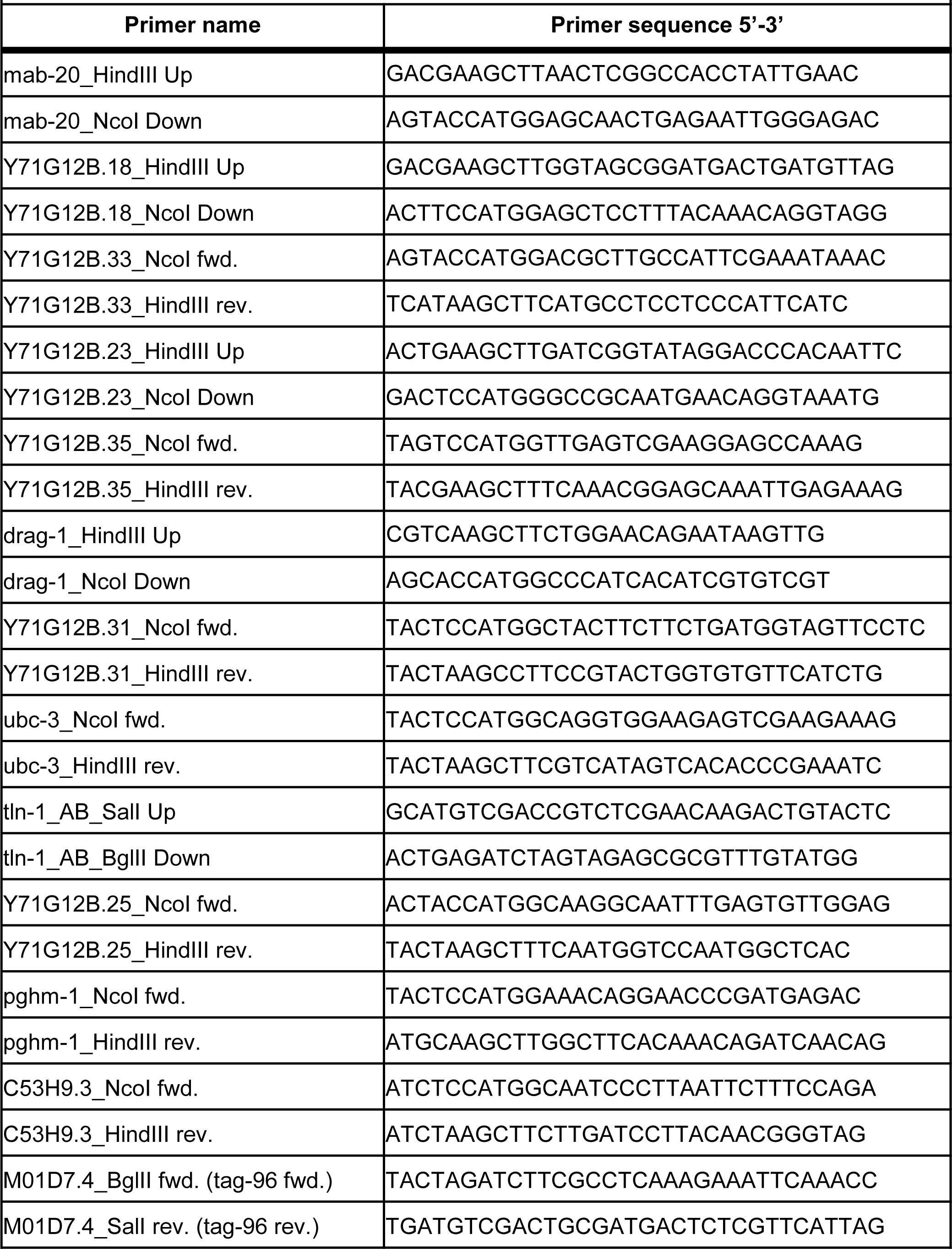
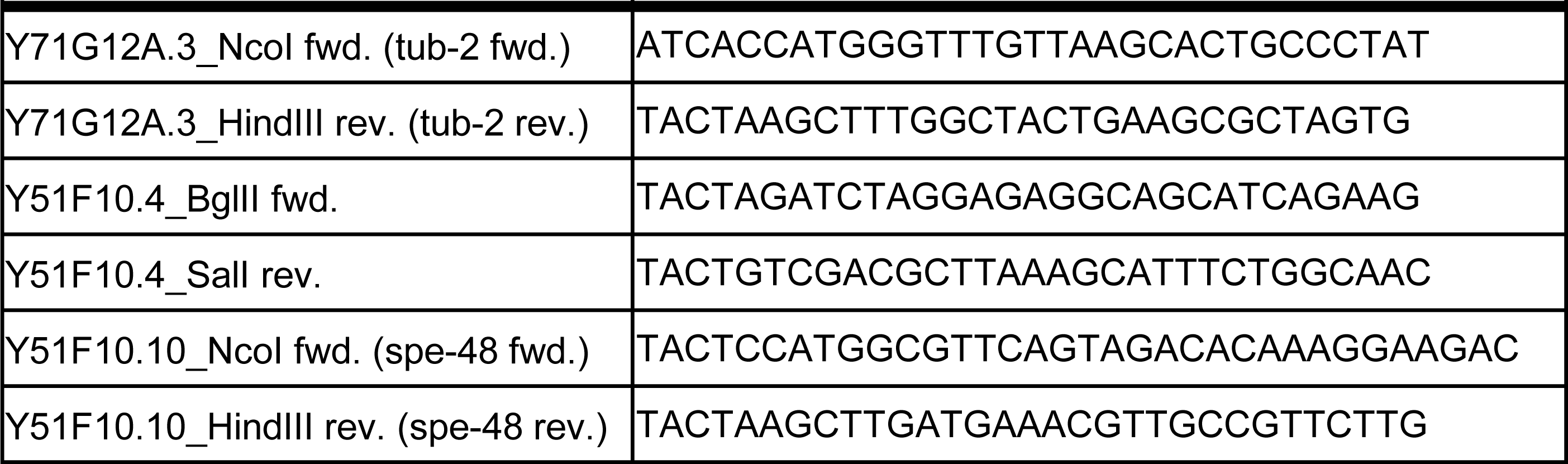
Primers used for generating RNAi clones.

**Data File 1.**
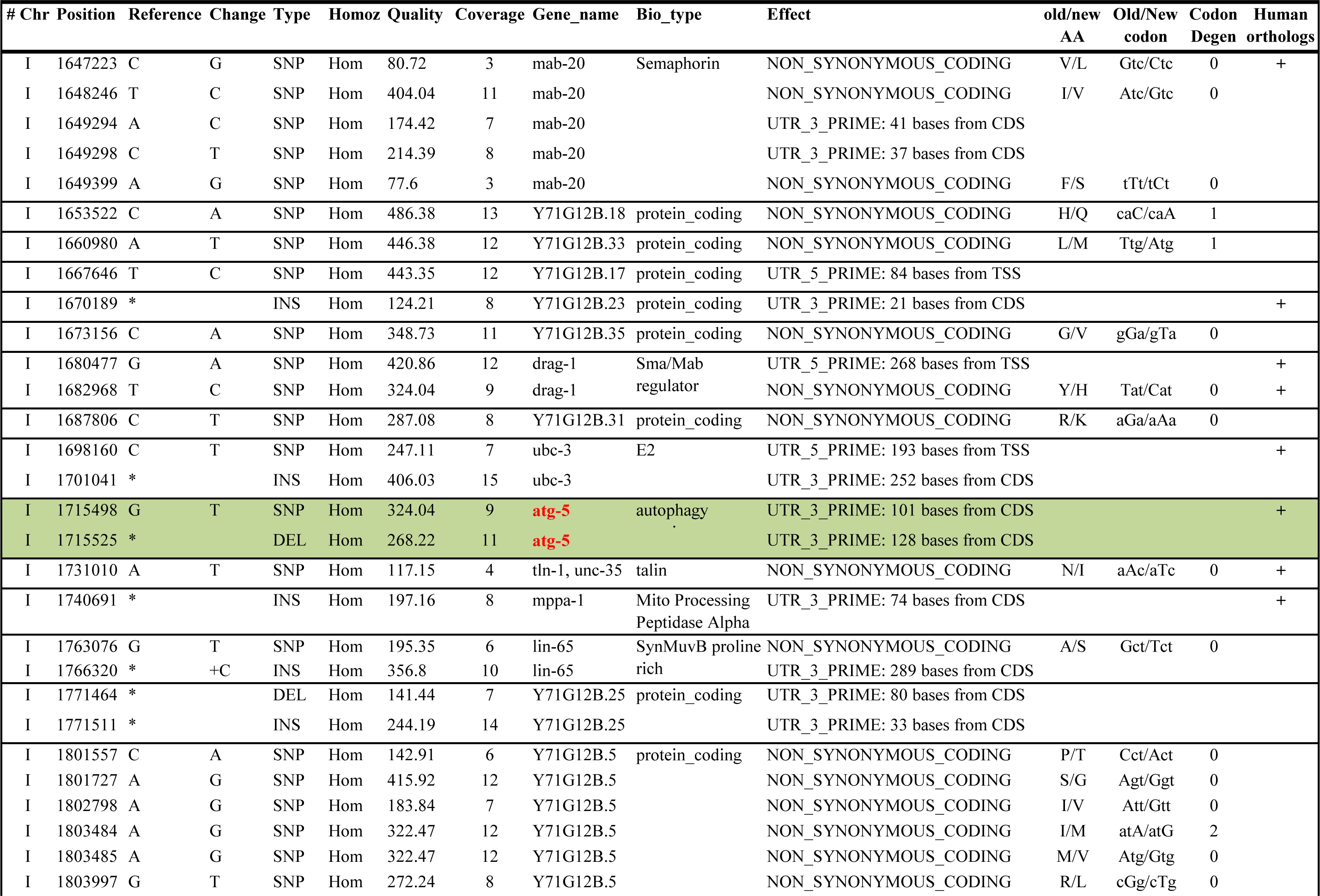
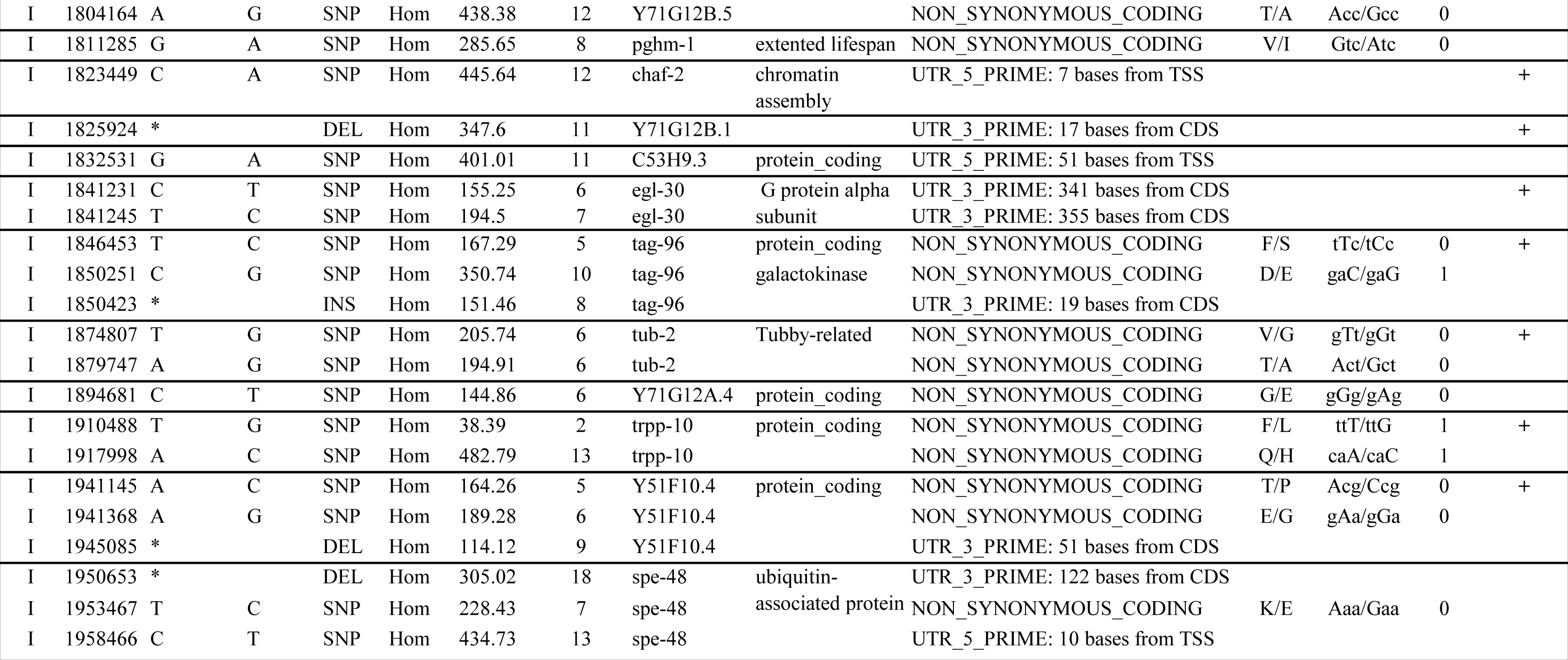
List of genes with potentially significant SNPs generated by the SnpEff tool. The nucleotide positions correspond to the N2(Bristol) genome assembly from WormBase release WS220 (122), available in the UCSC Genome Browser as ce10. The presence of human orthologs is according to (123).

